# Restoring glucose homeostasis with stomach-derived human insulin-secreting organoids

**DOI:** 10.1101/2022.12.15.520488

**Authors:** Xiaofeng Huang, Wei Gu, Jiaoyue Zhang, Ying Lan, Jonathan L. Colarusso, Sanlan Li, Christoph Pertl, Jiaqi Lu, Hyunkee Kim, Jian Zhu, Jean Sévigny, Qiao Zhou

**Affiliations:** Division of Regenerative Medicine & Ansary Stem Cell Institute, Department of Medicine, Weill Cornell Medicine, 1300 York Avenue, New York, NY 10065, USA; Département de Microbiologie-Infectiologie et d’immunologie, Faculté de Médecine, Université Laval, Québec City, QC G1V 0A6, Canada

## Abstract

Gut stem cells are accessible by biopsy and propagate robustly in culture, offering an invaluable resource for autologous cell therapies. Insulin-producing cells can be induced in mouse gut, but it has not been possible to generate abundant and durable insulin-secreting cells from human gut tissues to evaluate their potential as a cell therapy for diabetes. Here we describe a protocol to differentiate cultured human gastric stem cells (hGSCs) into pancreatic islet-like organoids containing gastric insulin-secreting (GINS) cells that resemble β-cells in molecular hallmarks and function. Sequential activation of the inducing factors NGN3 and PDX1-MAFA led hGSCs onto a novel differentiation path, including a SOX4^High^ endocrine and Galanin^High^ GINS precursor, before adopting β-cell identity, at efficiencies close to 70%. GINS organoids acquired glucose-stimulated insulin secretion in 10 days and restored glucose homeostasis for over 100 days in diabetic mice after transplantation, providing proof of concept for a new approach to treat diabetes.

---

Gut stem cells are highly proliferative and power the weekly self-renewal of the gut mucosal lining^1–3^. Harvested from biopsies, human gut stem cells can be propagated in culture as organoids or primary cell lines over many generations, providing abundant tissues for potential autologous transplantation therapies^4–6^. Gut stem cells produce gut-specific tissues, including hormone-secreting enteroendocrine cells (EECs). Rare insulin expressing EECs have been reported in fetal human small intestine^7^. Whether such cells secret insulin is unknown, but their presence suggests an intrinsic permissiveness for insulin production in the fetal if not postnatal intestine. Prior to this discovery, it was shown that suppressing FoxO1 could activate insulin in murine intestinal EECs^8^ and that pancreatic endocrine-like cells could be generated from putative human endocrine progenitors^9^. We also reported that co-expression of the endocrine regulator NEUROG3 (also known as NGN3) and pancreatic β-cell regulators PDX1 and MAFA could induce insulin-secreting cells from murine intestine and stomach^10,11^. However, the same approaches yielded few insulin producers from human gut organoids^11,12^.

Generating functional insulin-secreting cells has tremendous therapeutic value, offering treatments for insulin-dependent diabetes, including the autoimmune type 1 diabetes^13–18^. An attractive feature of using gut stem cells to make β-cell mimics is the ease of establishing autologous organoids from biopsies, which can enable mass production and personalized therapies. Aside from the current technical inability to differentiate gut stem cells into functional β-like cells at sufficient efficiency, a significant unknown factor is the documented short lifespans of gut cells in vivo, numbering in days to several weeks^19,20^. This raises the concern as to whether insulin-secreting cells made from human gut tissues will be sufficiently stable and durable as an engraftable therapeutic.

In this study, we developed a robust 3-step protocol to induce cultured hGSCs from multiple donors to differentiate into islet-like organoids at high efficiency, containing approximately 70% β-like cells and other islet-like endocrine populations. GINS organoids exhibited glucose responsiveness 10 days after induction. They were stable upon transplantation for as long as we tracked them (6 months), secreted human insulin and reversed diabetes in mice. No proliferative cells were detected in transplanted GINS organoids whereas hGSCs perished upon engraftment. GINS organoids thus possess favorable attributes as a potential transplantable therapeutic. We further characterized the novel differentiation path leading from hGSCs to GINS organoids with single cell RNA-sequencing (scRNA-seq). This study establishes a promising approach to procuring autologous human insulin producers for diabetes treatment.

## Results

### Generating islet-like GINS organoids from human stomach samples

Our previous study in mice suggested that stomach tissues are more amenable to adopting β-cell fate than intestinal tissues^10^. We therefore focused on using human gastric stem cells (hGSCs) to generate insulin-secreting cells. The human stomach has three distinct parts: the corpus, pylorus (antrum), and cardia, with corpus mucosa being most abundant. Biopsy samples from all three regions have been grown successfully as organoids in three-dimensional (3D) Matrigel or as 2D flat stem cell colonies, while maintaining their regional identity in culture^9,21,22^. After in vitro differentiation, hGSCs produce gastric mucosal cells including acid- and mucus-secreting cells^9,21,22^. In this study, we primarily used corpus tissues because of its ready availability.

For the ease of scaling stem cell production, we used 2D culture to expand hGSCs. Each biopsy-sized gastric sample typically yielded 30-40 primary colonies, which can be amplified to > 10^9^ cells within 2 months (Extended Data Fig. 1a, b). Cultured hGSCs continued to express the stomach stem/progenitor marker SOX9 and the proliferative marker KI67 after many passages (Extended Data Fig. 1a), consistent with prior report^9^. To explore ways to direct hGSCs into functional insulin secretors, we began with the NPM factors (*Ngn3, Pdx1*, and *Mafa*). This combination has been shown to confer varying degrees of β-cell properties to non-β cells including pancreatic acinar cells, duct cells, and others^23–27^. However, coexpression of the NPM factors using our previously published polycistronic cassette yielded low insulin expression in cultured hGSCs (Extended Data Fig. 1c). We therefore systematically evaluated conditions that may influence GINS cell formation including the timing of NPM expression, inclusion of additional genetic factors, and medium composition. Several notable observations include: (1) high-level insulin induction required transient NGN3 (for 2 days) followed by stable PDX1 and MAFA expression. This sequence of transgene activation was superior to NPM co-expression or expressing PDX1-MAFA prior to NGN3 (Extended Data Fig. 1d); (2) inclusion of additional β-cell fate regulators such as MAFB or NKX6-1 did not enhance insulin activation (Extended Data Fig. 1e). Critically, we formulated a fully chemically defined serum-free medium for GINS cell differentiation. From a screen of 23 supplements, some of which are employed in the induction of β-like cells from pluripotent stem cells, we found that Nicotinamide and Y-27632 (a ROCK inhibitor) significantly promoted *INS* mRNA levels whereas A8301 (an ALK5 inhibitor) stimulated spontaneous aggregation of nascent GINS cells and expression of several key β-cell transcription factors (Extended Data Fig. 2). The final GINS differentiation medium contained the three supplements, N2, B27, and N-acetyl cysteine in the basal Advanced DMEM/F12 medium.

For inducible NGN3 activation, a *Ngn3* and estrogen receptor (*ER*) fusion gene (*Ngn3ER*) was incorporated into the hGSCs by lentivirus^28^. hGSC differentiation was initiated by 4OH-Tamoxifen treatment of cultured *Ngn3ER-*hGSCs for two days (step 1), followed by lentiviral integration of a *Pdx1-Mafa* co-expression cassette (over 95% infection rate; step 2). Four days later, the nascent GINS cells were aggregated into spherical organoids (step 3), which can persist in the defined medium for up to four weeks (Fig. 1a, b, Extended Data Fig. 3a). Immunohistochemistry of GINS organoids from multiple donors revealed that majority of the organoid cells expressed c-peptide (CPPT, 65.4% ± 5.2%), whereas a minority expressed either glucagon (GCG, 2.2% ± 1.3%), somatostatin (SST, 6.1% ± 2.7%), or ghrelin (GHRL, 4.0% ± 1.5%) (Fig. 1c, d, Extended Data Fig. 3b). The cellular composition of organoids was further evaluated by flow cytometry, with the fractions of CPPT^+^, SST^+^ and GCG^+^ cells largely concordant with the immunostaining data (Fig. 1e). Using a cocktail of SST, GCG and GHRL antibodies, we determined that approximately 92.6% of all CPPT^+^ cells (or 61.0% of all organoid cells) were mono-hormonal (Extended Data Fig. 3c). The average insulin mRNA levels and insulin content of GINS organoids were comparable to primary islets (Fig. 1f, Extended Data Fig. 3d). GINS organoids from multiple donors expressed key β-cell markers *ABCC8, KCNJ11, GCK, PAX6*, and *NKX2-2*, at comparable levels to primary islets (Extended Data Fig. 3e).

**Fig. 1.**
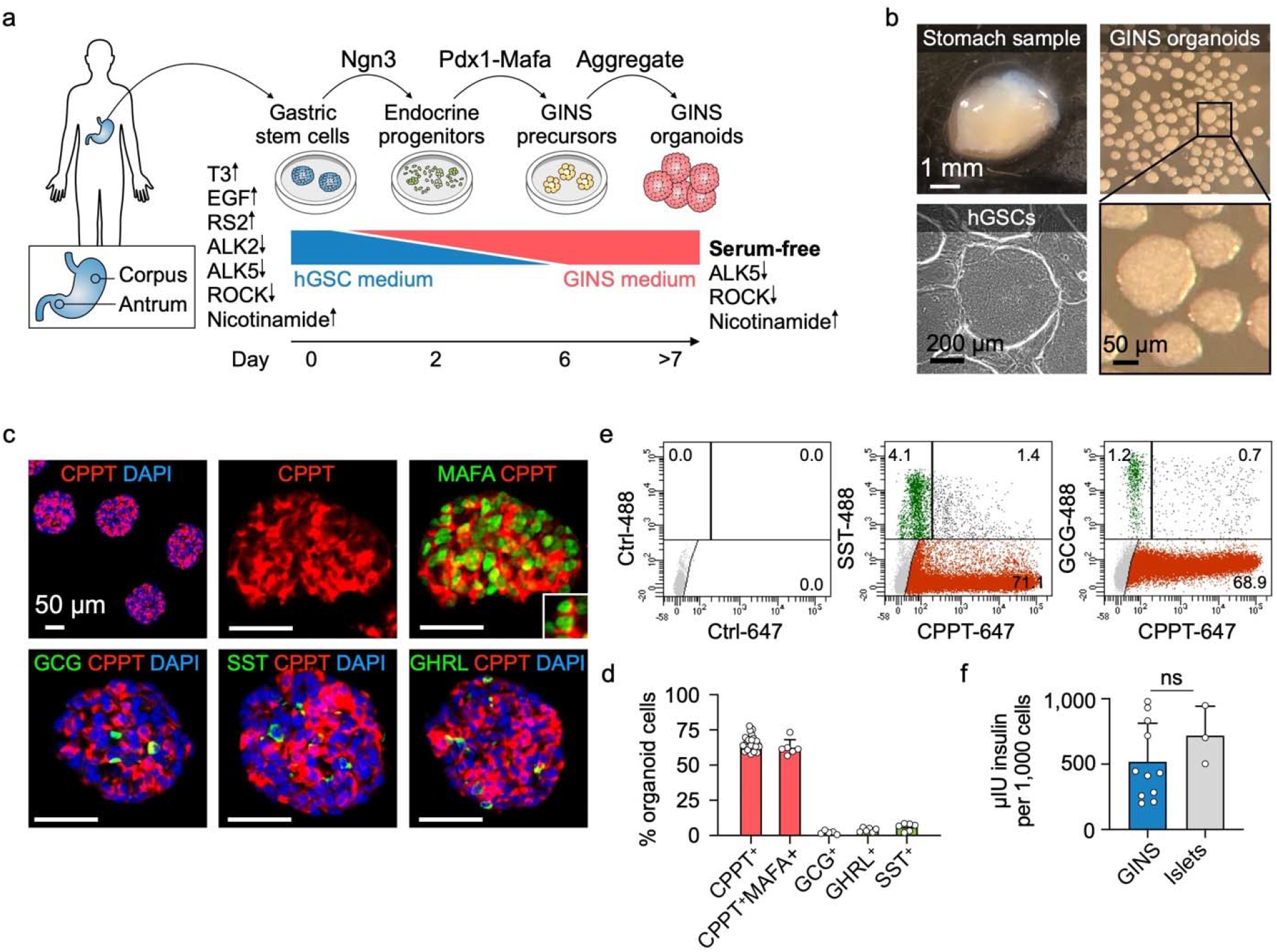
Generation of Gastric Insulin-Secreting (GINS) organoids from human stomach samples. **a,** Diagram showing key steps of GINS organoid generation. Medium supplements for hGSC (human gastric stem cell) culture or GINS organoid induction are indicated, with up and down arrows indicating agonists or antagonists, respectively. **b,** Representative images of human stomach samples, hGSC colonies, and GINS organoids. 500 cells per organoids. **c, d,** Immunofluorescent staining and quantification of day-21 GINS organoids for C-peptide (CPPT), MAFA, glucagon (GCG), somatostatin (SST), and ghrelin (GHRL). n = 3-6 separate batches of organoids from donor #6. **e,** Quantitative flow cytometry of day-21 GINS organoids (donor #10) with negative control. **f,** Insulin content of day-18 GINS organoids (n = 11 different batches of organoids from donor #6) and human islets (n = 3 samples from independent donors). Data are mean ± s.d.; *P* = 0.3006 (ns), unpaired t-test.

GINS organoids acquired glucose-stimulated insulin secretion (GSIS) 8-10 days after differentiation (Fig. 2a). Notably, glucose responsive organoids could be produced from multiple donors and maintained a batch to batch consistency in functionality (Fig. 2a, Extended Data Fig. 3f). However, the glucose responsiveness of GINS organoids became less consistent after prolonged culture (over 21 days) (Extended data Fig. 3f). GINS organoids responded to repeated glucose challenges as well as the clinical anti-diabetic drug Glibenclamide and the anti-hypoglycemia drug Diazoxide (Fig. 2b, Extended Data Fig. 3g). In dynamic GSIS assays, GINS cells from two separate donors responded robustly to KCl and liraglutide, a GLP-1 analog, but less so to high glucose challenge (Fig. 2c), indicating that they have not attained full functional maturity. Altogether, these data establish a GINS differentiation protocol robust for different donor tissues, yielding glucose-responsive organoids at high efficiency.

**Fig. 2.**
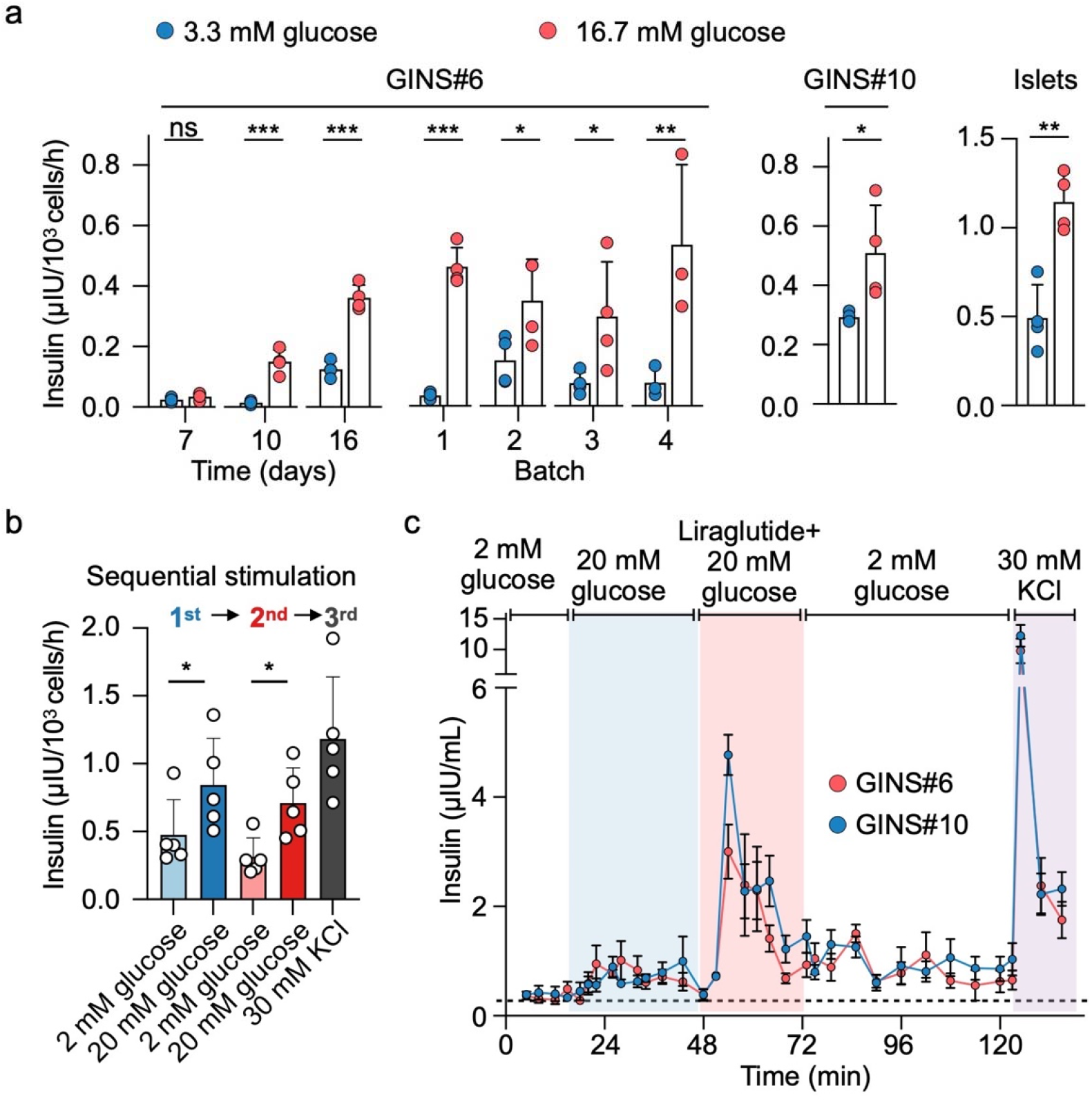
GINS organoids secret human insulin in response to glucose and GLP-1 analogue. **a,** Glucose-stimulated insulin secretion of GINS organoids at different time points (days post differentiation, 4 groups of same batch organoids per time point), from different batches (n=3-4 groups of organoids per batch) or from a different donor (n = 4 separate batches, day 21). \**P* < 0.05, \*\**p* < 0.01, \*\*\**p* < 0.001. For GINS organoids from donor #6, two-way repeated-measures ANOVA with Holm–Sidak’s multiple comparisons test. For GINS organoids from donor #10 and islets, one-tailed paired t-test. **b,** Insulin secretion of day-18 GINS organoids in response to sequential glucose challenges and KCl depolarization (n = 5 groups of organoids from donor #6, from two independent experiments). **c,** Dynamic glucose-stimulated insulin secretion by perifusion assay of day-21 GINS organoids derived from donor #6 and #10. Data presented as mean ± s.d. (n = 4 independent groups of organoids for each donor).

### GINS organoids contain four endocrine cell types

To better understand the identities of the cells in GINS organoids, we used scRNA-seq to interrogate the transcriptomes of 6,242 organoid cells (Fig. 3a). Clustering with published scRNA transcriptomes of human islets revealed four endocrine cell types that aligned with islet β-, α-, δ-, or ε-cells^29^ (Fig. 3a). No pancreatic polypeptide-positive cells were detected in GINS organoids (Fig. 3a). Clustering with hGSCs and mucus-secreting cells (derived from spontaneous hGSC differentiation in culture) showed almost no gastric cells remaining in GINS organoids (Extended Data Fig. 4a, 4b). The α and δ-like endocrine cells expressed canonical markers of their islet counterparts including *GCG, ARX, TTR*, and *GC* in α-cells, and *SST* and *HHEX* in δ-cells (Fig. 3b, Extended Data Fig. 4b). GINS cells expressed classical human β-cell markers including *G6PC2, GCK, ABCC8, NKX2-2, PCSK1, PAX6*, and key genes involved in β-cell identity, metabolism, insulin synthesis and secretion, and ion channel activities, but did not express *NKX6-1* (Fig. 3c, Extended Data Fig. 4c). Several genes previously shown to interfere with proper glucose sensing, including *HK1, LDHA*, and *SLC16A1*^30^, were strongly down-regulated in GINS cells (Extended Data Fig. 4d). GINS organoid cells ceased proliferation after differentiation (Extended Data Fig. 4e). Single-cell transcriptomes of antral GINS organoids also revealed a dominant fraction of β-like cells. No significant numbers of α- and ε-like cells were detected whereas an additional gastrin-expressing cell population was present in antral organoids (Extended Data Fig. 5a-e). Antral organoids exhibited robust GSIS in vitro (Extended Data Fig. 5f).

**Fig. 3.**
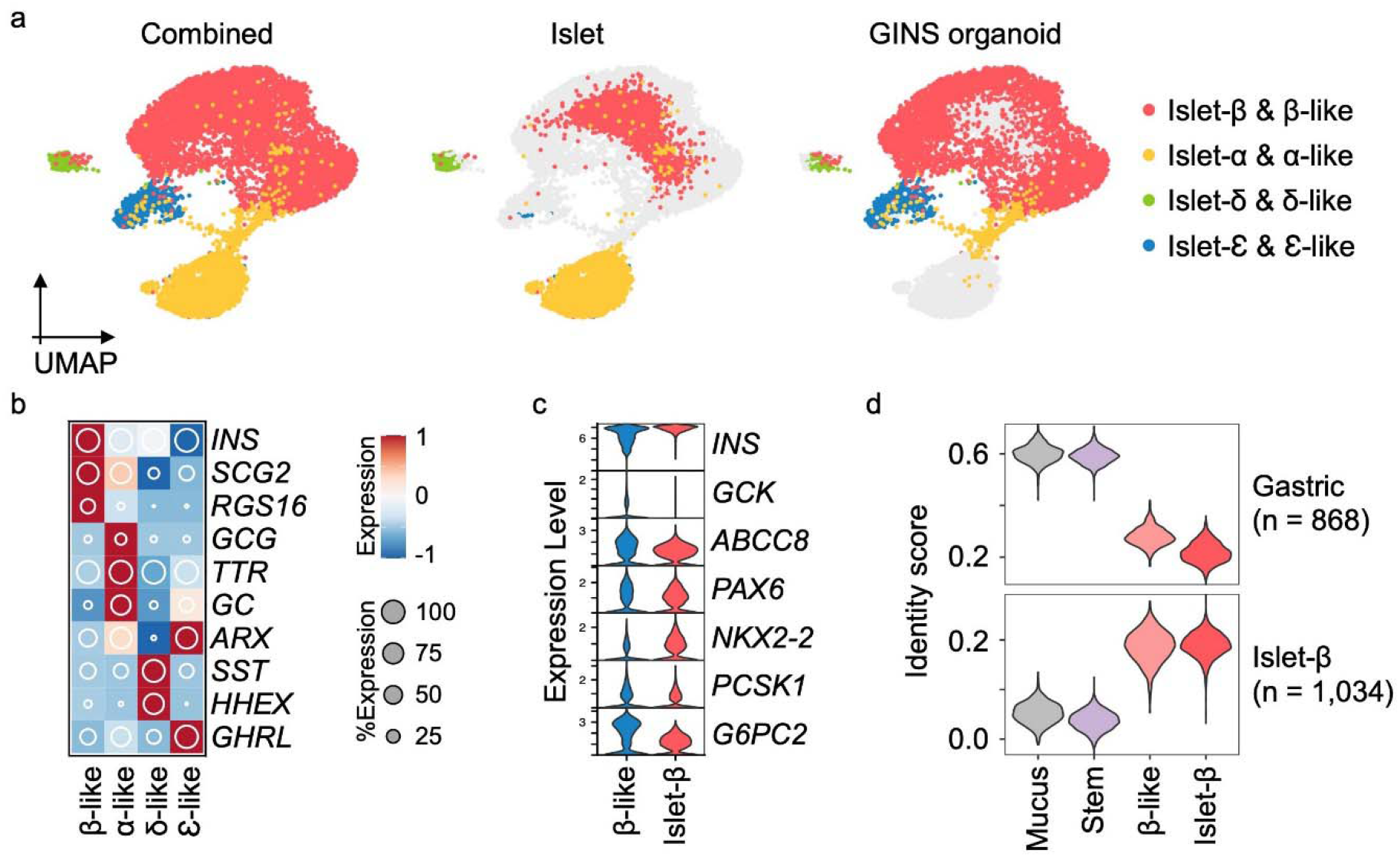
Four endocrine cell types identified in GINS organoids by scRNA-seq. **a,** UMAP visualization of integrated GINS organoid (day 21, donor #6) and primary human islet single-cell transcriptomes. A pancreatic polypeptide-positive cell population present in islets but not GINS organoids is not shown. **b,** Relative expression of endocrine cell type-specific markers. The shading displays scaled average gene expression, and diameter denotes fractional expression. **c,** Violin plots of key β-cell function and identity markers in GINS β-like and islet β cells. **d,** Cell identity scoring with gastric and β-cell signatures (868 and 1,034 genes, respectively). Mucus: mucus-secreting cells spontaneously differentiated from hGSCs in culture; Stem: hGSCs; β-like: GINS β-like cells; islet-β: islet β-cells.

To further assess the identity of GINS cells, we applied molecular score cards of β-cells (1,034 β-cell-specific genes) and gastric cells (868 stomach-specific genes) benchmarked from published human scRNA data^31,32^. GINS cells scored similarly to islet β-cells in both categories, although there is a residual gastric signature in GINS cells (Fig. 3d). In comparison, hGSCs and mucus-secreting cells possessed low β- and high gastric scores (Fig. 3d). These data indicate that GINS cells possess the general molecular identity of islet β-cells at the single-cell level, consistent with their glucose responsiveness. Nevertheless, Gene Ontology analysis suggests that GINS cells have lower ribonucleoprotein biogenesis activity than islet β-cells (Extended Data Fig. 6), possibly underlying the functional immaturity of GINS cells in vitro.

### GINS cells persist after transplantation and reverse diabetes

In order to evaluate the longevity and functionality of GINS cells in vivo, we transplanted GINS cells under the kidney capsule of immune-compromised NSG mice (0.8 million cells per mouse). We examined the grafts at 2, 4, and 6 months. At each time point, the grafts contained abundant INS^+^ cells perfused with CD31^+^ vasculature and minor populations of GCG^+^, SST^+^, and GHRL^+^ cells (Fig. 4a, Extended Data Fig. 7a). Grafted GINS cells expressed PAX6, NKX2-2, PCSK1, and the adult β-cell marker ENTPD3^33^ (Fig. 4a). Electron microscopy showed that the electron-dense granules of the GINS cells were not fully condensed (Extended Data Fig. 7b), likely reflecting lower levels of SLC30A8 (Extended Data Fig. 7c), the activity of which is required for the granule morphology^34^. Loss of SLC30A8 is associated with protection against type 2 diabetes^35,36^.

**Fig. 4.**
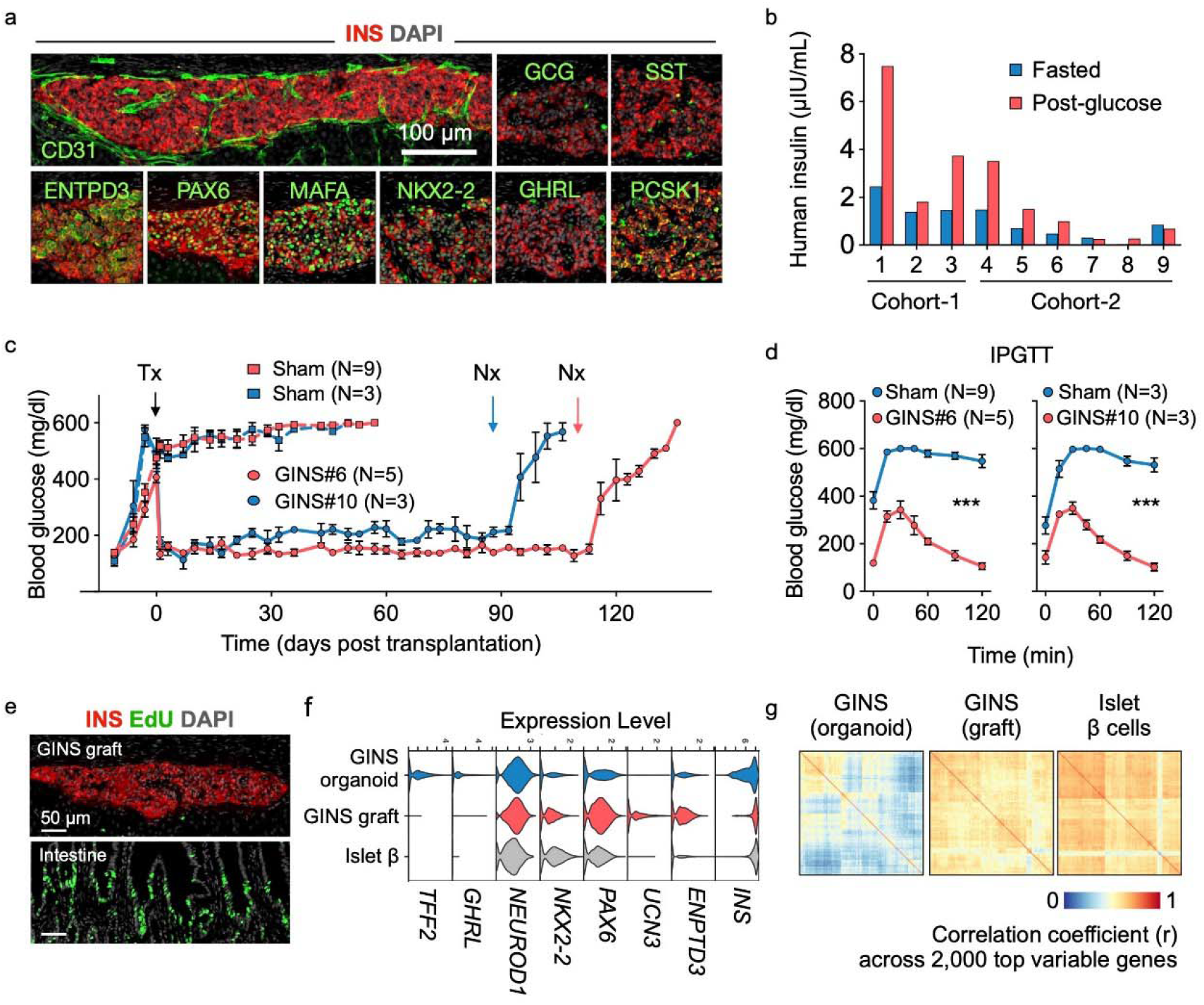
Transplanted GINS organoids secrete human insulin and reversed diabetes in mice. **a, b,** GINS cells (0.8 × 10^6^, corpus donor #6) were transplanted under the renal capsule of non-diabetic NSG mice, which yielded grafts containing CD31^+^ vascular cells and predominant INS^+^ cells that coexpressed PAX6, MAFA, NKX2-2, PCSK1, and the mature β cell-associated marker ENTPD3 (6 months post transplantation). A small number of SST^+^, GCG^+^, and GHRL^+^were also found **(a).**Human insulin from individual mouse was measured at 7 weeks (cohort-1) and 11 weeks (cohort-2) post transplantation, grafted with different batches of cells, after overnight fasting (blue bar) and 60 minutes after glucose challenge (red bar) **(b). c, d,** STZ-induced diabetic NSG mice were transplanted with 6-8 million GINS cells from donor #6 (GINS#6) or 6 million GINS cells from donor #10 (GINS#10) or without transplantation (Sham). Random-fed blood glucose levels **(c)** and intraperitoneal glucose tolerance tests (IPGTT, 2 weeks post transplantation) **(d)** showed significant improvement for both transplanted groups. Data presented as mean ± s.e.m..; *P* < 0.001 (***), two-way repeated-measures ANOVA. Tx: transplantation. Nx: nephrectomy. **e**, 24-hour pulse-chase with EdU revealed no proliferative cells in GINS graft but abundant replicating cells in the intestine of the same engrafted mouse. **f**, Violin plots of select genes in GINS β-like cells before and after transplantation and comparison with islet β cells. **g**, Assessment of transcriptomic heterogeneity among GINS β-like cells before and after transplantation and among primary human islet β cells with correlation coefficient analysis across 2,000 top variable genes.

The majority of the GINS grafts showed glucose-stimulated insulin secretion (Fig. 4b). Accordingly, transplantation of 6-8 million GINS cells from donor #6 into NSG mice rendered diabetic by streptozotocin (STZ) rapidly suppressed hyperglycemia and maintained glucose homeostasis for over 100 days, until removal of the grafts by nephrectomy (Fig. 4c, Extended data Fig. 7d). A second cohort of mice transplanted with organoids from a different donor (# 10, 6 million cells per mouse) yielded similar results although the glycemic control was less tight (Fig. 4c). Glucose tolerance improved significantly in both engrafted groups (Fig. 4d). Importantly, we detected no proliferating cells within the grafts at any time point (Fig. 4e). To directly evaluate the fate of hGSCs upon transplantation, we engrafted 0.5 million undifferentiated mCherry-labeled hGSCs under the kidney capsule of 6 NSG mice. After 80 days, no surviving cells were detected at the graft sites (Extended Data Fig. 7e), consistent with the well-documented reliance of hGSCs on high WNT signaling to survive^37,38^. We conclude that GINS cells, derived from human gut stem cells, can be long-lived and functional and pose little risk of uncontrolled proliferation after transplantation.

Comparison of GINS single-cell transcriptomes before and after transplantation (6,242 cells in vitro, 3,502 cells in vivo at 3 months post transplantation) showed molecular changes consistent with maturation, including enhanced expression of *NKX2-2, PAX6, UCN3*, and *ENTPD3* and reduced *TFF2* and *GHRL* (Fig. 3f). Key ribonucleoproteins were up-regulated whereas several pathways elevated in cultured GINS cells were down-regulated after transplantation (Extended Data Fig. 7f-h). Insulin expression, while variable in cultured GINS cells, became notably more uniform in the grafted cells (Fig. 4f). Correlation coefficient analysis based on the top 2,000 variable genes showed that GINS transcriptomes became more homogeneous after transplantation, a characteristic shared with islet β cells (Fig. 4g). These results together indicate molecular maturation of GINS cells after transplantation.

### Developmental trajectory of GINS cells

hGSCs normally produce stomach-specific cells including mucus- and acid-secreting cells. To understand how their differentiation path is rerouted in GINS formation, we used scRNA-seq to sample key stages in GINS derivation and reconstructed the developmental trajectory with pseudotime ordering. In total, 9,544 high-quality single cell transcriptomes were collected from four samples: hGSCs, endocrine progenitors (2 days after NGN3 activation), GINS precursors (4 days after PDX1-MAFA expression), and GINS organoids (14 days after aggregation) (Fig. 5a). Clustering analysis showed one stem cell (*SOX9^High^TFF2^High^LGR5^High^*), two endocrine progenitors (*SOX4^High^CHGA^Low^* and *SOX4^High^CHGA^High^*), one GINS precursor (*GAL^High^SSTR2^High^*), and four endocrine cell populations (Fig. 5b, c, Extended Data Fig. 8a) along pseudotemporal progression (Fig. 5d, Extended Data Fig. 8b). Notably, somatostatin-expressing δ-like cells emerged ahead of the other endocrine cells (Fig. 5a, 5b, 5d). *Pdx1* and *MafA* transgene expression were significantly higher in GINS cells than the other endocrine cells, suggesting higher transgenes promoted β-cell fate at the expense of the other endocrine cell types (Fig. 5e). Gene Ontology analysis showed rapid down-regulation of stem cell and proliferative pathways upon endocrine differentiation (Fig. 5f). WNT and NOTCH signaling were active in endocrine precursors whereas histone modification was associated with the GINS precursors. Functional pathways characteristic of β-cells such as hormone transport and secretion, and mitochondria and ribosome activities, emerged last (Fig. 5f).

**Fig. 5.**
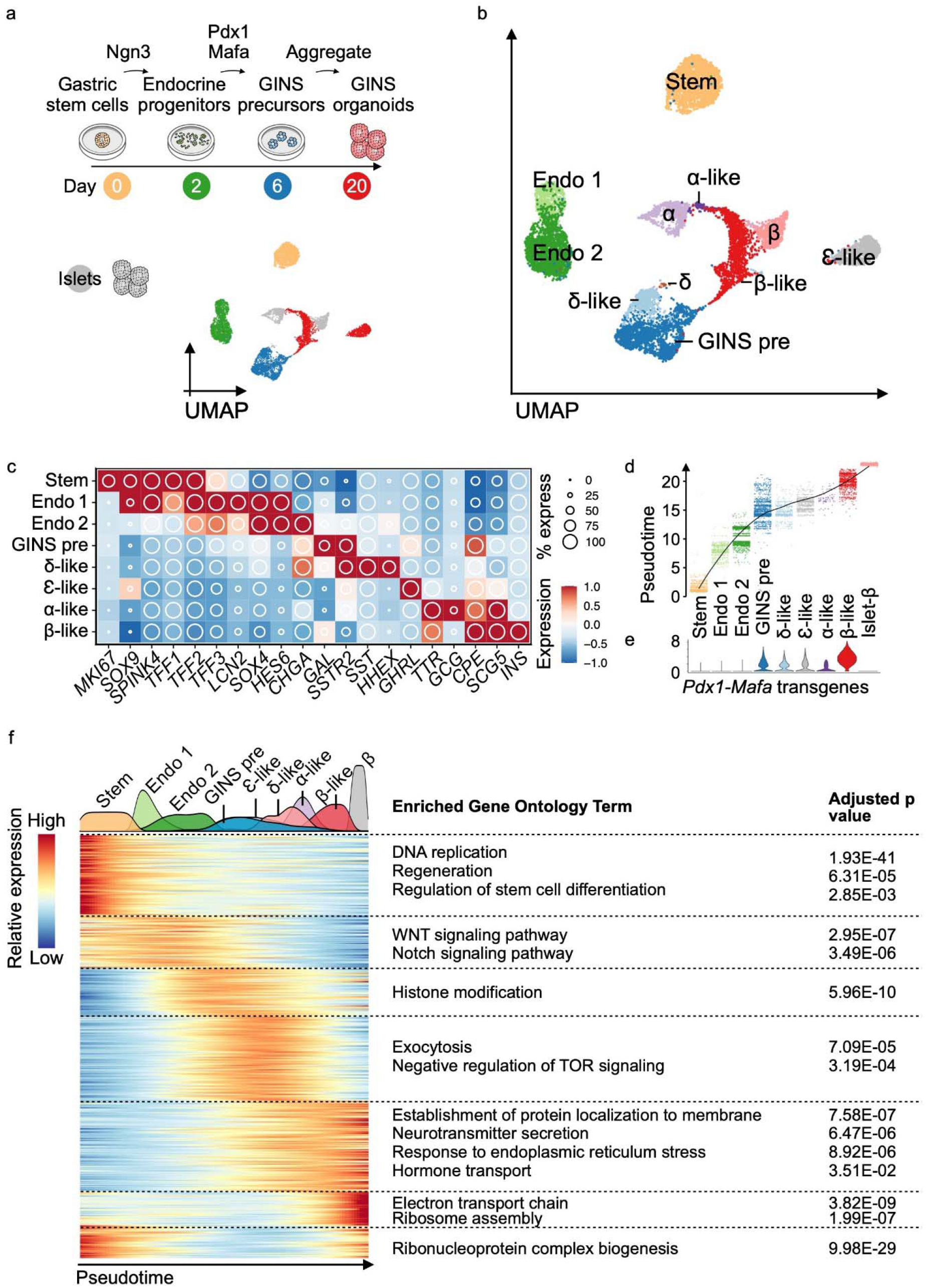
Developmental trajectory of GINS cells. **a,** Experimental design of sampling cells with scRNA-seq at key stages of hGSC differentiation to GINS organoids. UMAP showing clustering of hGSCs (day *0*, yellow), endocrine progenitors (day 2, green), GINS precursors (day 6, blue), GINS organoids (day 20, red) and primary human islets (gray). **b,** UMAP of cell types colored according to their identities. Stem: hGSCs; Endo 1: endocrine progenitors 1; Endo 2: endocrine progenitors 2; GINS pre: GINS precursors. **c,** Relative expression of select markers across cell types. **d,** Pseudotime trajectory analysis of cell types. **e,** Violin plot showing the expression levels of the *Pdx1-Mafa* transgenes in different cell types. **f,** Heat map showing gene-expression clusters along the pseudotime trajectory from hGSCs to GINS cells and human islet β cells. Select significant GO terms enriched in each gene cluster are shown.

Waves of transcription factor (TF) activations accompanied the progression from hGSCs to GINS cells, likely orchestrating the stepwise acquisition of GINS fate (Extended Data Fig. 8c). Regulon analysis showed active ASCL1 and SOX4 regulons in endocrine progenitors (Extended Data Fig. 9a, 9b). Early-activating GINS regulons included ones for HHEX, PAX4, and ISL1, while late-activating regulons included ones for RFX6, PAX6, PDX1 and MAFB (Extended Data Fig. 9a, 9b).

Pseudotime ordering and RNA velocity analysis suggest that β-like cells descended from GINS precursors (Fig. 5d, Extended Data Fig. 9c), which expressed several markers including *SSTR2* and *GALANIN* (GAL), a neuropeptide predominantly expressed in the nervous system (Fig. 6a). To evaluate whether GAL^+^precursors can give rise to CPPT^+^ cells, we made a reporter construct in which GFP expression was driven by human *GAL* promoter and integrated this construct into a hGSC line (Fig. 6b). Six days post differentiation, GFP expression was activated, consistent with the appearance of GINS precursors at this stage. We then purified the GFP^high^ cell fraction by flow cytometry and aggregated the cells into organoids. After overnight culture, the nascent GINS organoids contained predominant GAL^+^ cells (89.9% ± 3.0%) whereas a small fraction (14.2% ± 5.9%) had barely detectable levels of CPPT (Fig. 6c-e). After 14 days of culture, the percentage of GAL^+^ cells and the average GAL staining intensity decreased while the percentage of CPPT^+^ cells rose to 61.2% ± 7.5% and CPPT staining intensity significantly increased (Fig. 6c-e). These data show that GAL^+^ cells can indeed serve as precursors to CPPT^+^ GINS cells. We note that GALANIN is expressed in human islets, including some β-cells (Fig. 6f). Upon transplantation and maturation, *GAL* expression in GINS cells further decreased (Fig. 6g). Altogether, our data support a model in which sequential Ngn3 and Pdx1-Mafa expression triggered waves of TF activations that led gut cells onto a novel differentiation path, including a galanin-expressing precursor, before adopting GINS identity (Fig. 6h).

**Fig. 6.**
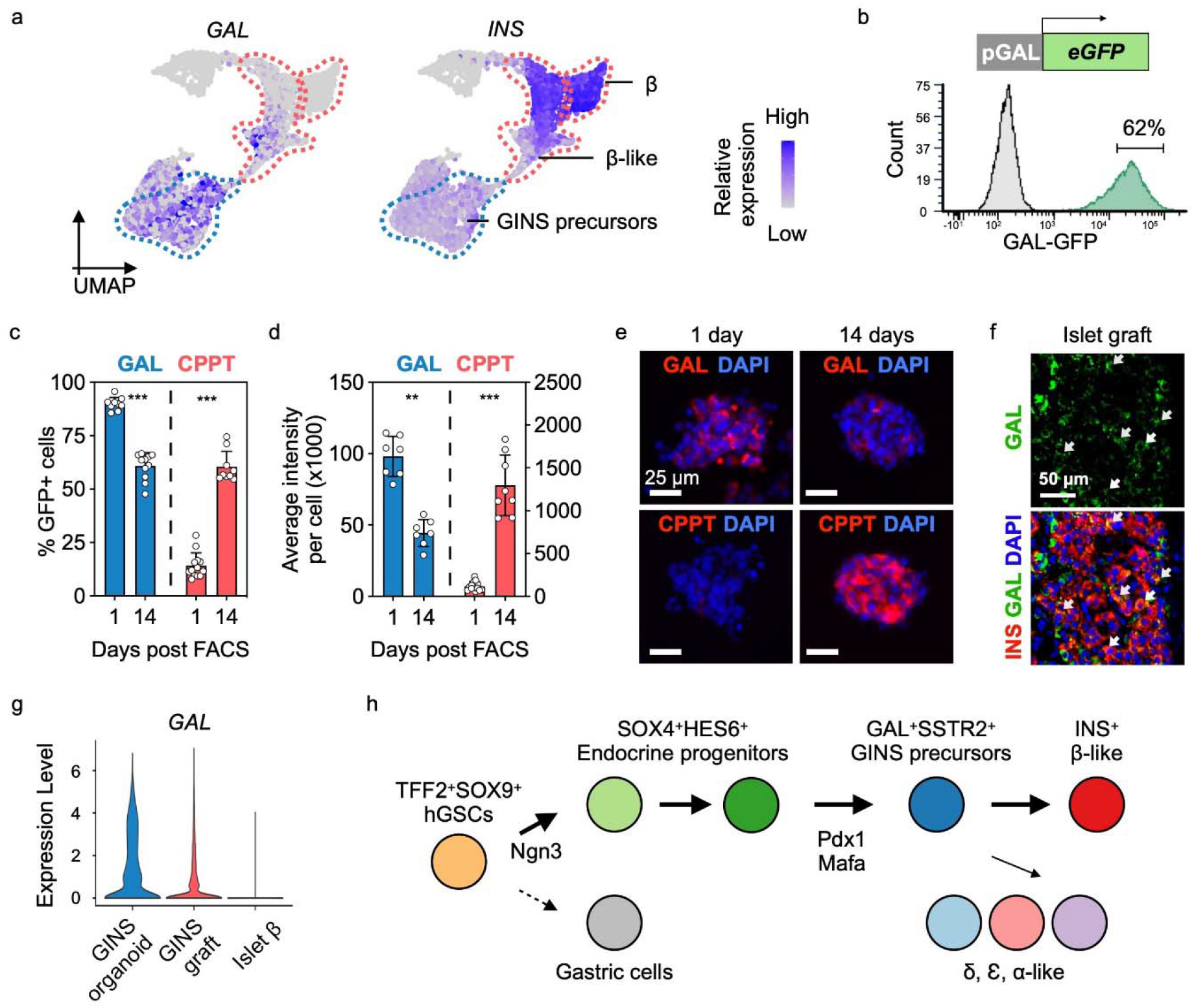
Galanin^+^ precursors give rise to GINS cells. **a,** Relative expression of *INS* and *GAL* in GINS precursors, GINS β-like cells and human islet β cells. UMAP subset of Fig. 5b. **b,** Purification of GFP^High^ GINS precursors differentiated from *GAL-GFP* hGSC reporter line at day 7 post differentiation. **c, d,** Quantification of GAL^+^ and CPPT^+^ cell percentage in organoids and their average staining intensity at day 1 or 14 post sorting. (n=7-12 GINS organoids from donor #6). **e,** Representative images of immunofluorescent staining for GAL and CPPT at day 1 and day 14 post sorting. **f,** Representative immunofluorescence of GAL and INS in primary human islet grafts in NSG mice. Arrows indicate GAL^+^INS^+^ cells. **g,** Violin plot showing scRNA levels of *GAL* in GINS β-like cells before and after transplantation and in human islet β cells. **h,** Proposed model for GINS organoid formation, with rerouting of the hGSC developmental trajectory by NGN3, PDX1 and MAFA.

## Discussion

In this study, we describe a robust protocol to derive insulin-secreting organoids from cultured hGSCs with high efficiency. The glucose-responsive GINS cells became long-lived insulin secretors and restored glucose homeostasis in diabetic mice. Some of the attractive features of using gastric stem cells to make β-cell mimics include: the relative ease of obtaining gastric biopsies via endoscopy to enable production of autologous grafts; GINS organoid generation involves a relatively straightforward 3-step protocol; GINS organoids gain a measure of functionality relatively quickly, at 10 days post induction; GINS organoids pose little tumorigenic risks; the same protocol works well on hGSCs from different donors; and lastly, GINS organoids persist for many months after transplantation.

The leading strategy of producing islet-like organoids today is differentiation from human embryonic stem cells, yielding sc-islets^39^. An autologous approach using induced pluripotent stem cells (iPSCs) is feasible but faces significant hurdles^18^, one of which is the lengthy process in generating iPSCs. The hGSC approach saves time and effort by directly obtaining patient stem cells and producing organoids with a simpler differentiation protocol, thus offering a useful alternative to iPSCs.

In addition to the autologous approach, more than 250,000 bariatric surgeries, in which a large portion of the gastric corpus is removed, are performed yearly in the United States alone. These discarded tissues could support the building of gastric stem cell biobanks, enabling major histocompatibility complex (MHC) matching to find suitable donors for transplantation. For treatment of type 1 diabetes, recent advances in engineering immune-evasive organoids could guide the generation of GINS organoids that survive an autoimmune environment

GINS grafts, unlike sc-islet grafts, lack a significant number of α- and δ-like cells. These endocrine populations have been suggested to fine-tune glucose responsiveness of β-cells^43^. The long-term consequence of this particular composition of GINS grafts requires further study and if necessary, a new method should be developed to boost the number of α- and δ-like cells.

Our data uncovered significant differences between mouse and human gastric tissues in their response to the inducing NPM factors. Whereas mouse corpus stomach resisted NPM induction^9^, human corpus cells were amenable. And while simultaneous expression of NPM factors induced abundant β-like cells from mouse antral EECs in situ^9^, it induced few from cultured human antrum and corpus stem cells. The molecular basis for these species-specific effects remains to be determined.

Although GINS cells resemble cadaveric β-cells, there are notable differences, including the retainment of a residual gastric molecular signature in GINS organoids. We formulated a fully chemically defined serum-free medium for GINS organoid differentiation, but the organoids did not attain full functionality in vitro and after prolonged culture, their glucose-responsiveness waned. Developing a new medium to promote organoid maturation in vitro will be a priority of future studies. Additional optimizations are also needed to move the GINS technology toward clinics, including replacement of lentivirus with a safer gene activation method.

---

**Extended Data Fig. 1.**
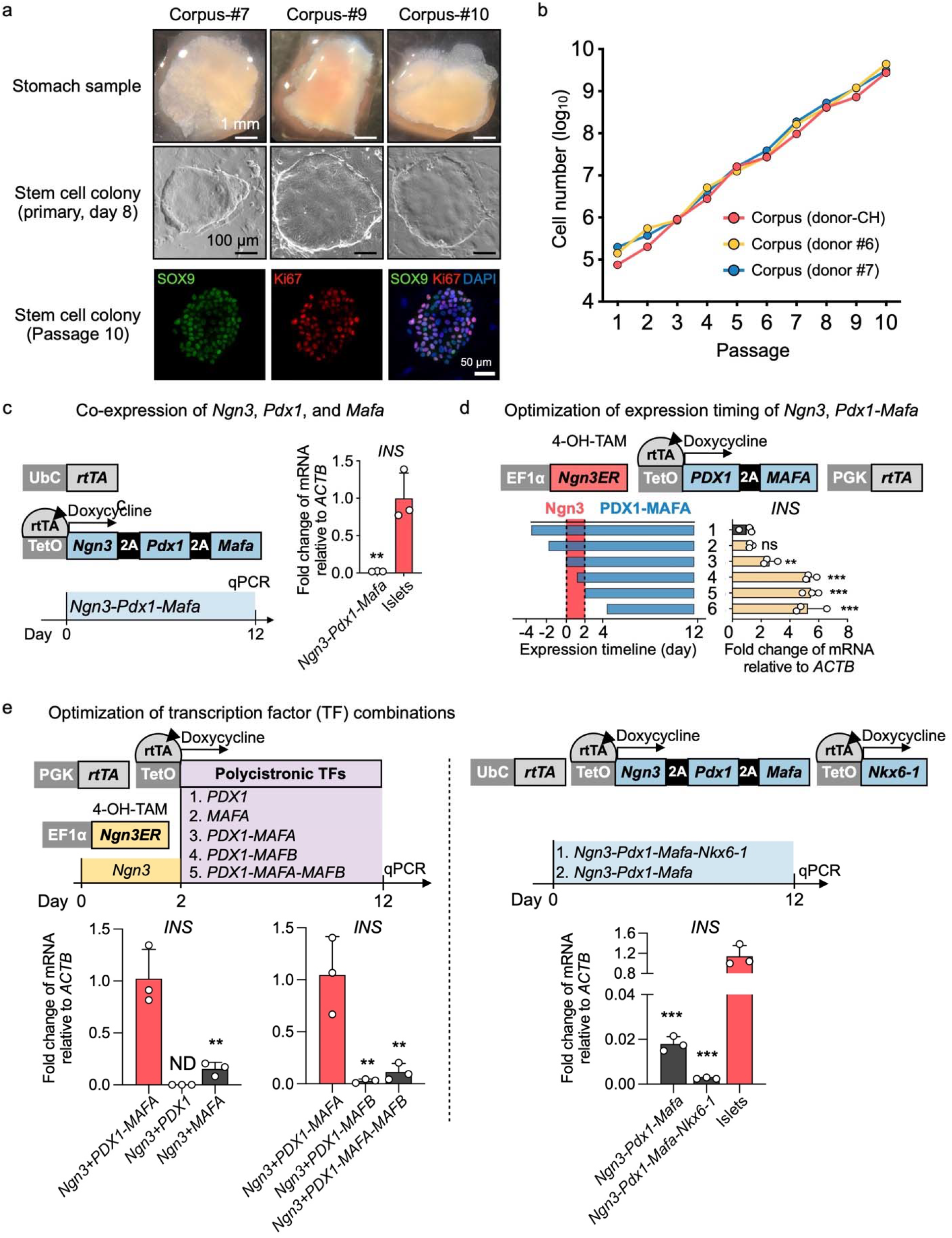
Optimizing conditions to induce insulin-expressing cells from cultured human gastric stem cells. **a,** Top: stomach samples from 3 different donors; middle, primary hGSC colonies derived from the stomach samples; bottom, immunofluorescence of passage-10 hGSC colony staining for SOX9 and KI67. **b,** Growth kinetics of hGSCs from three different donors. **c,** Co-expression of *Ngn3, Pdx1*, and *Mafa* using a polycistronic inducible construct in hGSCs yielded low levels of insulin expression (n=3 biological independent samples). Ubc: ubiquitin promoter. **d,** To optimize the relative timing of Ngn3 and Pdx1-MafA expression, we expressed a Ngn3ER fusion protein in which Ngn3 activity was induced by 4-OH Tamoxifen (4-OH-TAM). Polycistronic PDX1 and MAFA co-expression was controlled by rtTA-TetO and activated by the addition of Doxycycline in the culture medium. Higher *INS* expression was achieved by sequential activation of the transcription factors NGN3 and PDX1-MAFA. **e,** Comparison of PDX1-MAFA with the other transcription factor combinations in insulin induction. 2-day Ngn3ER induction (by 4-OH-TAM) preceded the other TFs, or alternatively, co-expression cassettes were used. ND: not detected. **c, d, e,** Data presented as mean ± s.d.; \**P* < 0.05, \*\**P* < 0.01, \*\*\**P* < 0.001, by two-tailed unpaired t-test **(c),** or one-way ANOVA with Dunnett multiple comparisons test **(d, e)**.

**Extended Data Fig. 2.**
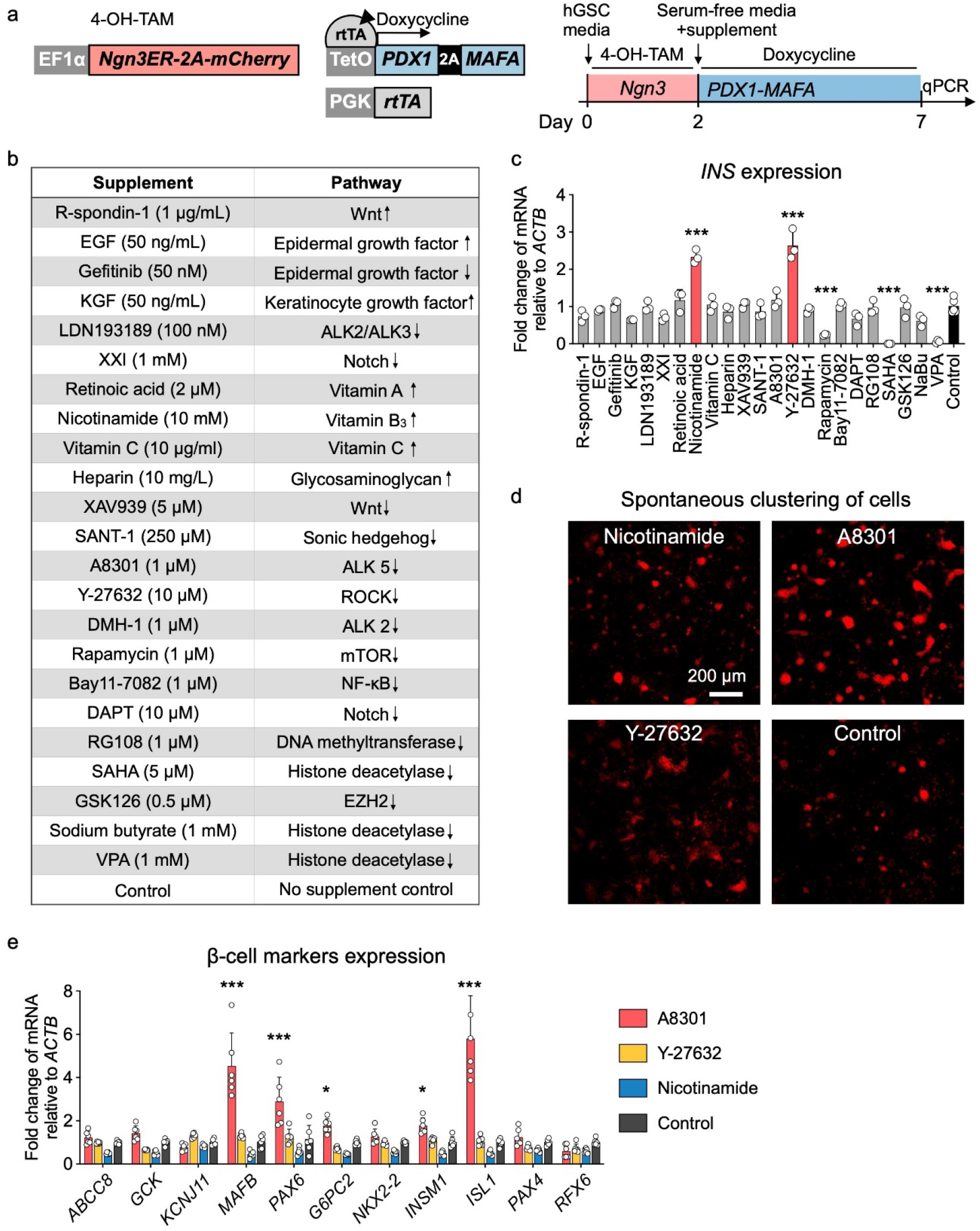
Formulating chemically defined serum-free medium for GINS organoid differentiation. **a,** Experimental design for the supplement screen. *Ngn3ER:* fusion gene in which NGN3 activity was induced by 4-OH-TAM. Polycistronic PDX1 and MAFA co-expression was controlled by rtTA-TetO and activated by the addition of Doxycycline in the culture medium. mCherry was co-expressed with Ngn3ER in the cell line. Ngn3 was activated from day 0 to day 2. Culture medium was switched to a basal serum-free medium (advanced DMEM/F12, 10 mM HEPES, 1X GlutaMAX, 1X B-27, 1X N-2, and 500 μM N-Acetyl-L-Cysteine) on day 2 with addition of a single supplement and Doxycycline. **b,** The list of supplements that were screened and the pathways they targeted. Up and down arrows indicate agonists or antagonists, respectively. **c,** Relative expression of *INS* mRNA on day-7 post differentiation in comparison with no supplement control (n = 3 or 5, independent samples). Nicotinamide and Y-27632 treatment significantly up-regulated *INS*. **d,** Spontaneous clustering of cells was evaluated by mCherry live imaging on day-7 post differentiation. Select conditions were shown. A8301 treatment had the most observable clustering effect on the nascent GINS cells. **e,** Relative expression of β-cell markers measured on day-7 post differentiation in comparison with no supplement control (n=6, independent samples). **c, e,** Data presented as mean ± s.d.; \**P* < 0.05, \*\**p* < 0.01, \*\*\**P* < 0.001, by one-way **(c),** or two-way ANOVA **(e)** with Holm–Sidak’s multiple comparisons test.

**Extended Data Fig. 3.**
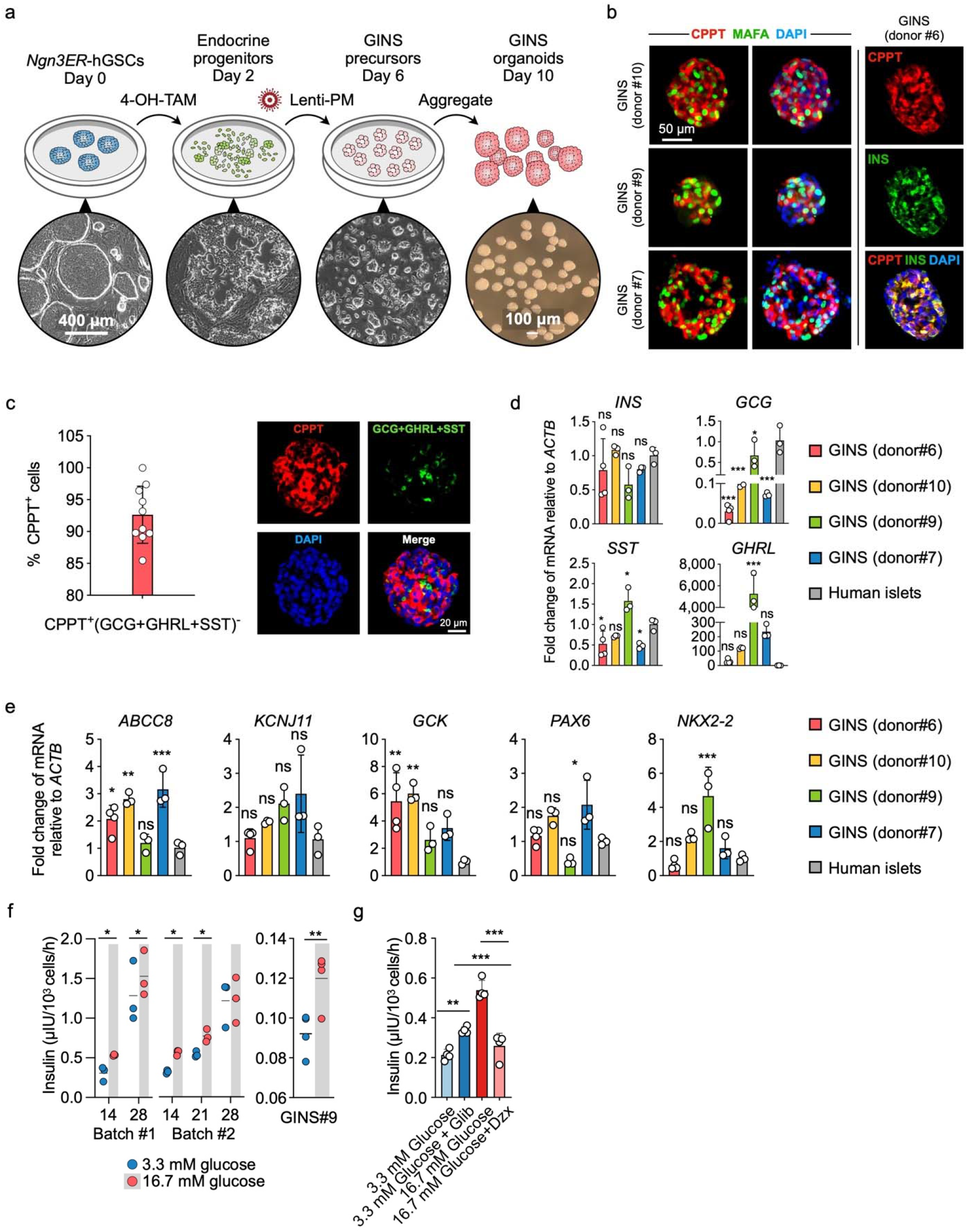
Molecular and functional characterization of GINS organoids derived from multiple donors. **a,** Schematic diagram and representative images of cells at key stages in GINS organoid formation. *Ngn3ER-hGSCs:* human gastric stem cells that incorporated a *Ngn3* and estrogen receptor (*ER*) fusion gene (*Ngn3ER*); 4-OH-TAM: 4-OH Tamoxifen; Lenti-PM, lentiviral integration of a polycistronic *Pdx1-Mafa* co-expression cassette. **b,** Representative immunofluorescent staining of corpus GINS organoids derived from three different donors, and co-localization of INS and CPPT in GINS cells. **c,** To assess CPPT^+^mono-hormonal cells, a cocktail of GCG, SST and GHRL antibodies were stained together with CPPT in day-21 GINS organoids. Right panel shows immunofluorescent staining of CPPT (red) and a combination of GCG, GHRL and SST (green), left panel shows quantification of mono-hormonal CPPT^+^ cells (n = 10 organoids from donor #6). **d,** Relative expression of endocrine hormone genes including *INS, GCG, SST*, and *GHRL* in day-18 GINS organoids derived from four different donors in comparison with human islets (n = 3-4 separate batches of samples for each donor). **e,** Relative expression of key β-cell markers in GINS organoids derived from different donors in comparison with human islets (n = 3-4 separate batches of samples for each donor). **f,** Glucose-stimulated insulin secretion of GINS organoids at different time points (n = 3 independent samples from donor #6 for each batch of differentiation) or donor #9 (n = 3 independent samples, day-18). **g,** Insulin secretion of day-18 GINS organoids from donor #6 incubated with the indicated concentrations of glucose with or without 10 nM glibenclamide (Glib) or 0.5 mM diazoxide (Dzx) as indicated (n = 4 independent samples). **d-g,** Data presented as mean ± s.d. \**P* < 0.05, \*\**P* < 0.01, \*\*\**p* < 0.001, by one-way ANOVA **(d, e, g),** repeated-measures Two-way ANOVA with Holm–Sidak’s multiple comparisons test for different time points **(f)** or one-tailed paired t-test for donor#9 **(f).**

**Extended Data Fig. 4.**
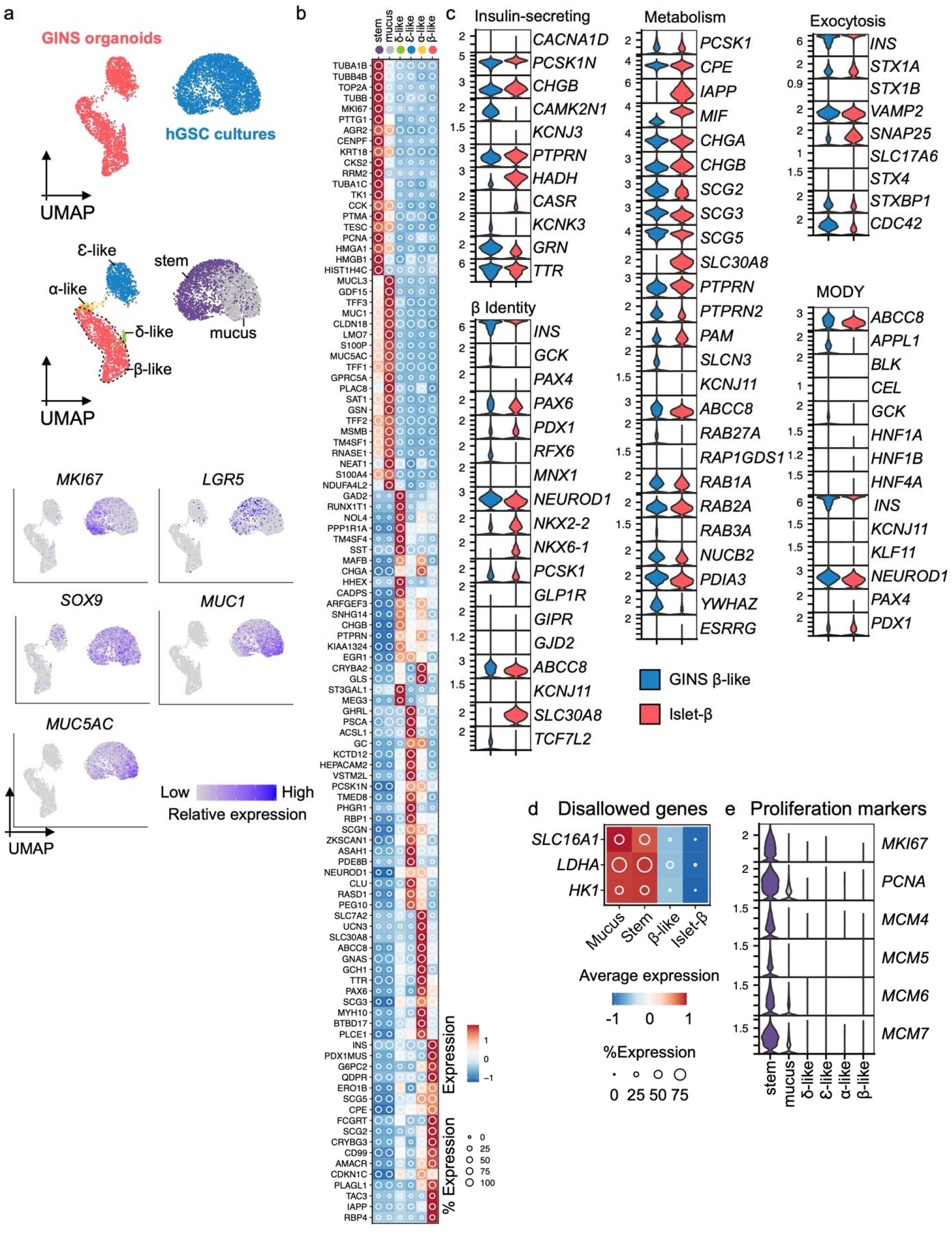
Characterizing GINS organoid cells with scRNA-seq. **a,** Top UMAP, cells sampled from hGSC cultures (blue) or GINS organoids (red); middle UMAP, hGSC cultures included both stem cells (stem) and mucus-secreting cells (mucus) spontaneously differentiated from hGSCs. GINS organoids contained four endocrine cell types. Cells are colored according to cell types; bottom UMAP, relative expression of cell type-specific markers. **b,** Relative expression of endocrine cell type-specific markers. The shading displays scaled average gene expression, and diameter denotes fractional expression. **c,** Comparison of GINS β-like cells and islet β-cells in expression profiles of key genes for β-cell function, identity, metabolism, and exocytosis. MODY: Maturity Onset Diabetes of the Young. **d,** Relative expression of disallowed genes in the indicated cell types. **e,** Violin plots showing the expression levels of proliferative markers in the indicated cell types.

**Extended Data Fig. 5.**
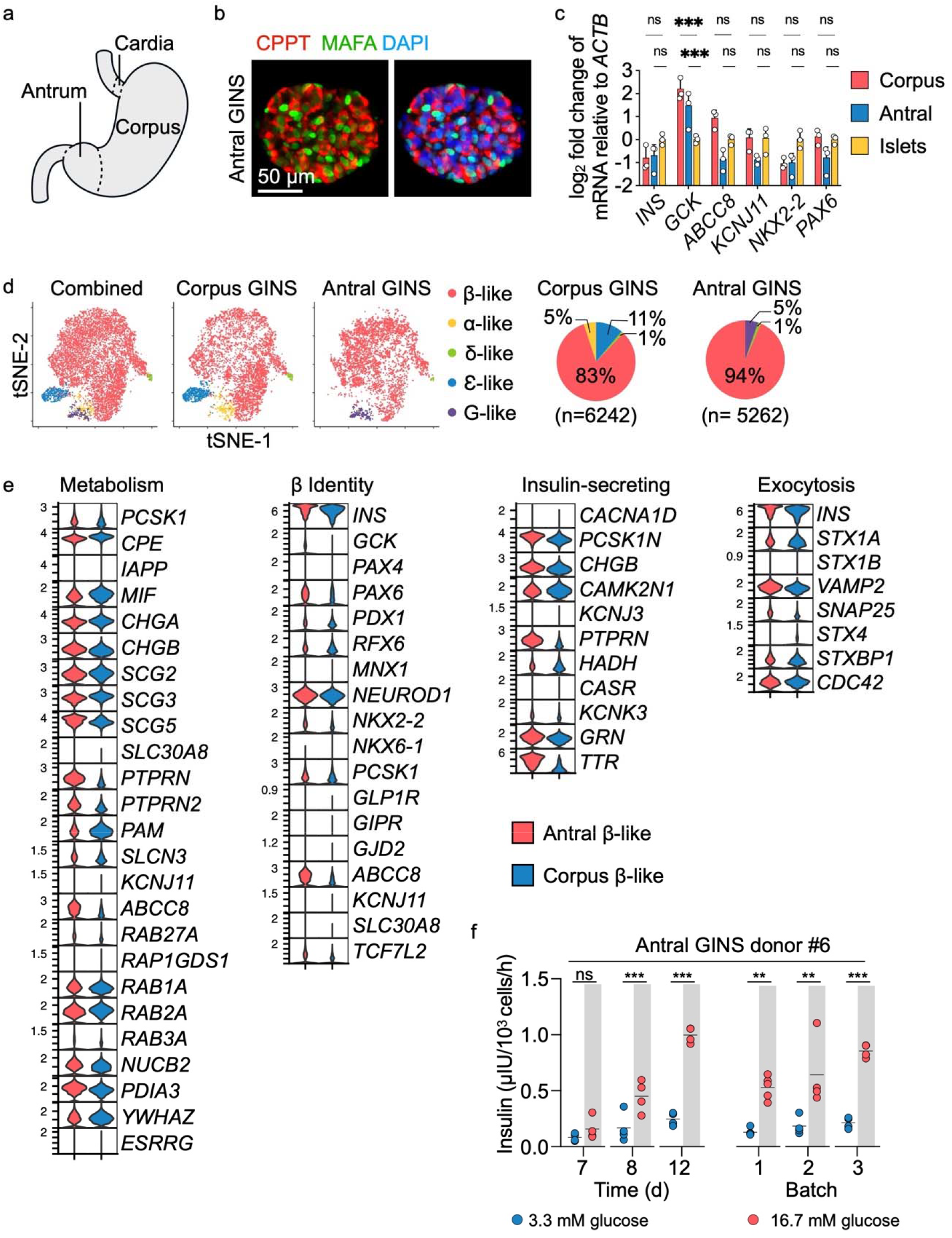
scRNA-seq comparison of GINS organoids derived from human antrum vs corpus stomach. **a,** Diagram of human stomach. **b,** Immunofluorescence of antral GINS organoid (donor#6) stained for CPPT and MAFA. *c*, Comparison of corpus and antral GINS organoids (both from donor#6) in the expression of β-cell marker genes. Data presented as mean ± s.d.; \*\*\**p* < 0.001, by one-way ANOVA with Dunnett multiple comparisons test comparing GINS with islets. **d,** t-distributed stochastic neighbor embedding (t-SNE) plots of integrated corpus and antral GINS organoids. Cells are colored according to cell types. G-like: G-like cells that expressed gastrin (*GAST*). Pie charts indicate cell-type proportions. **e,** Comparison of antral and corpus GINS β-like cells expression profiles of key genes for β-cell function and identity. Red, antral GINS β-like cells; Blue, corpus GINS β-like cells. **f,** Glucose-stimulated insulin secretion of antral GINS organoids at different time points (days post differentiation) or from different batches. \**P* < 0.05, \*\**p* < 0.01, \*\*\**p* < 0.001. Two-way repeated-measures ANOVA with Holm–Sidak’s multiple comparisons test.

**Extended Data Fig. 6.**
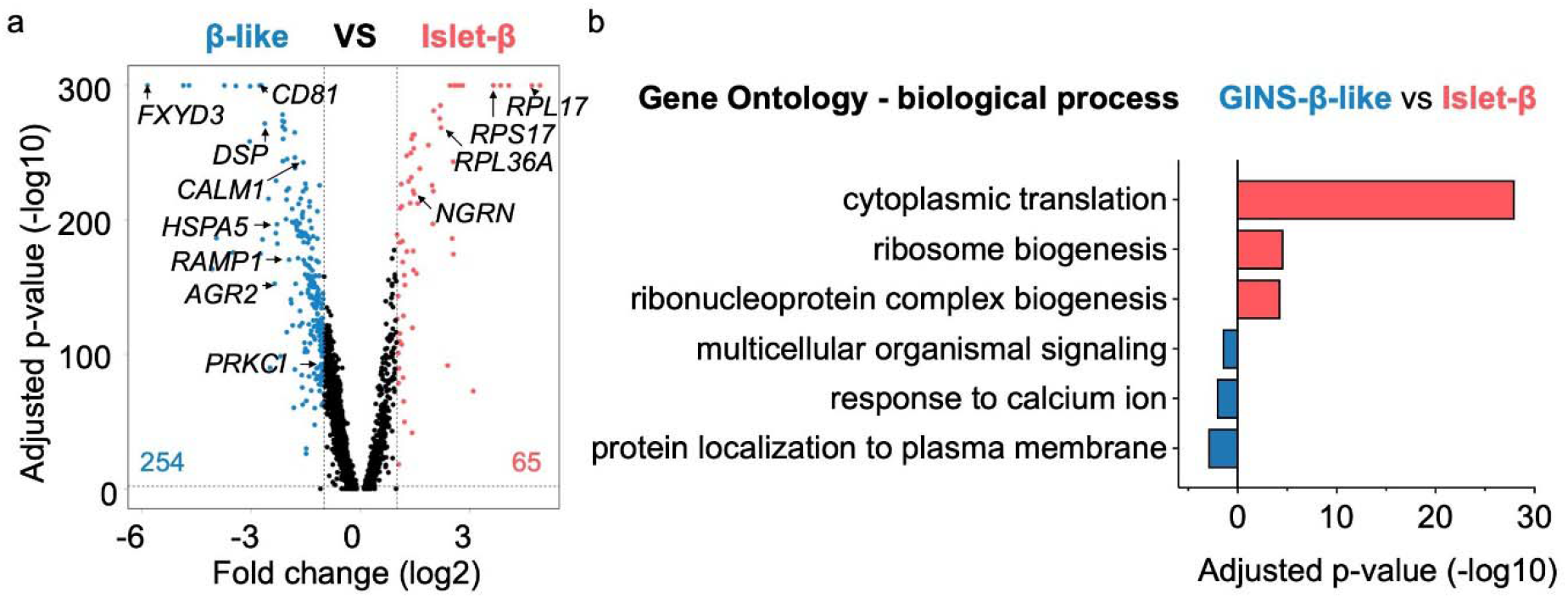
GINS organoids are not fully mature. **a,** Volcano plot comparing gene expression of GINS β-like cells versus islet β cells identified in Fig. 3a. The number of differentially expressed genes (DEGs) enriched in either cell group is shown in the plot. Threshold of DEGs: adjusted-P < 0.01 and log_2_ fold-change > 1. b, Gene Ontology (GO) analysis of DEGs enriched in GINS β-like cells (blue) or islet β-cells (red).

**Extended Data Fig. 7.**
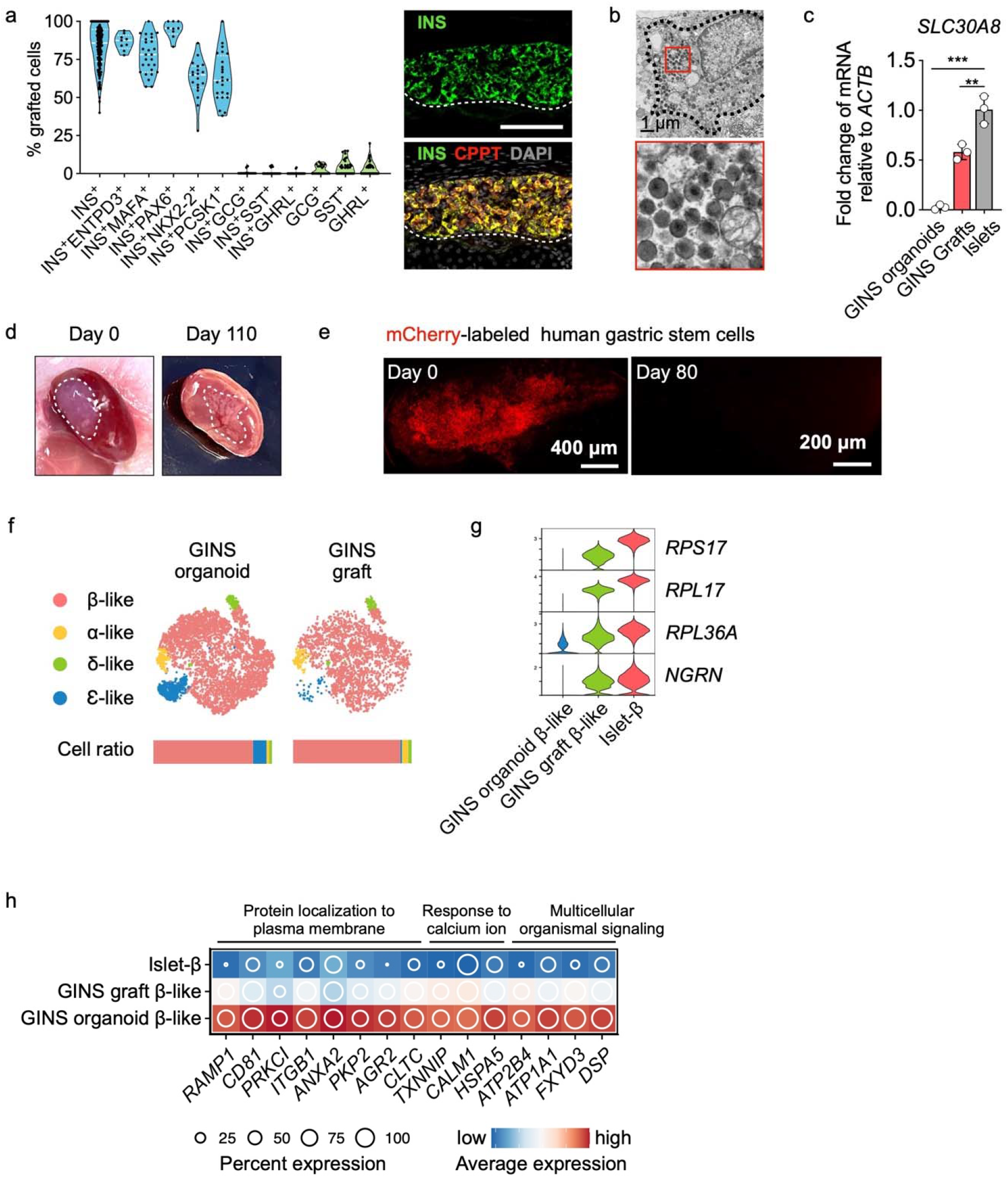
Phenotypic characterization of GINS grafts. **a,** Quantification from immunofluorescent staining of marker proteins, n = 3 independent experiments. Representative image showing co-expression of INS and CPPT in the GINS graft. **b,** Electron microscopy imaging of GINS graft. The electron-dense core granules were partially condensed. **c,** *SLC30A8* relative expression levels in GINS organoids, GINS grafts, and human islets. Data presented as mean ± s.d.; \*\**P* < 0.01, \*\*\**p* < 0.001, by one-way ANOVA with Dunnett multiple comparisons test comparing GINS with islets. **d,** Images of the kidney from mice transplanted with GINS cells on day 0 and day 110 post transplantation. **e,** mCherry-labeled hGSCs (0.5 × 10^6^) transplanted under the renal capsule and visualized under fluorescent microscope on day 0 and day 80 post transplantation. No Cherry^+^ cells were found at day 80. **f,** tSNE projection of integrated GINS organoids and grafts. Cells are colored according to cell types. Horizontal bars indicate cell type ratios. **g,** Violin plots showing the expression levels of select ribonucleoproteins. **h,** Relative expression of select genes in the pathways elevated in cultured GINS β-like cells compared with human islet-β cells.

**Extended Data Fig. 8.**
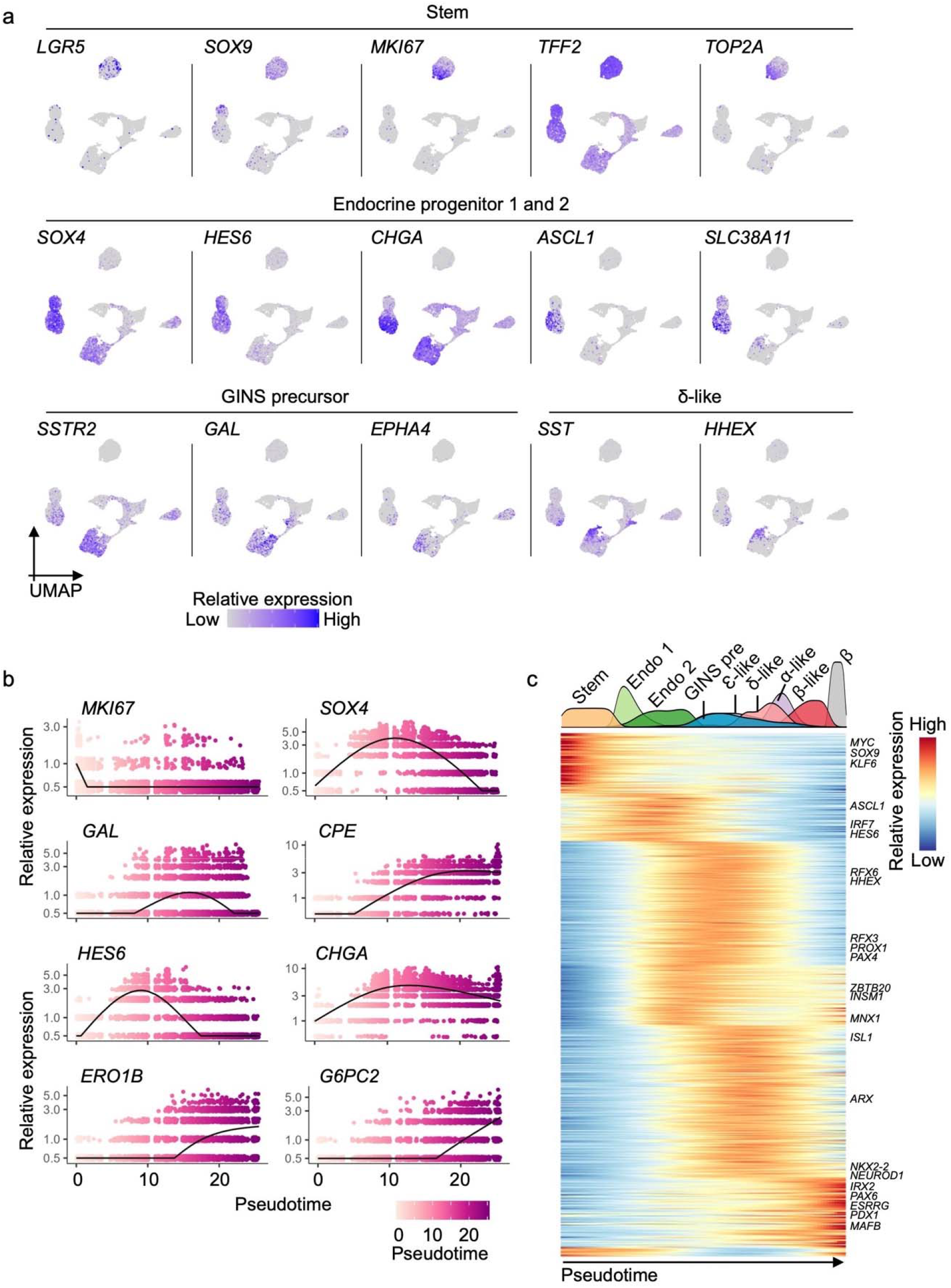
Dynamic gene and signaling pathway activations in hGSC differentiation to GINS organoid. **a,** Relative expression levels of cell type-specific markers in UMAP. **b,** Expression of select genes shown along pseudotime in GINS organoid formation. Each dot represents a cell. **c,** Heat map showing transcription factor expression clusters along the pseudotime trajectory from hGSCs to GINS organoid cells and islet β cells. Density plot on the top showing cell populations along pseudotime. Stem: hGSCs; Endo 1: endocrine progenitors 1; Endo 2: endocrine progenitors 2; GINS pre: GINS precursors.

**Extended Data Fig. 9.**
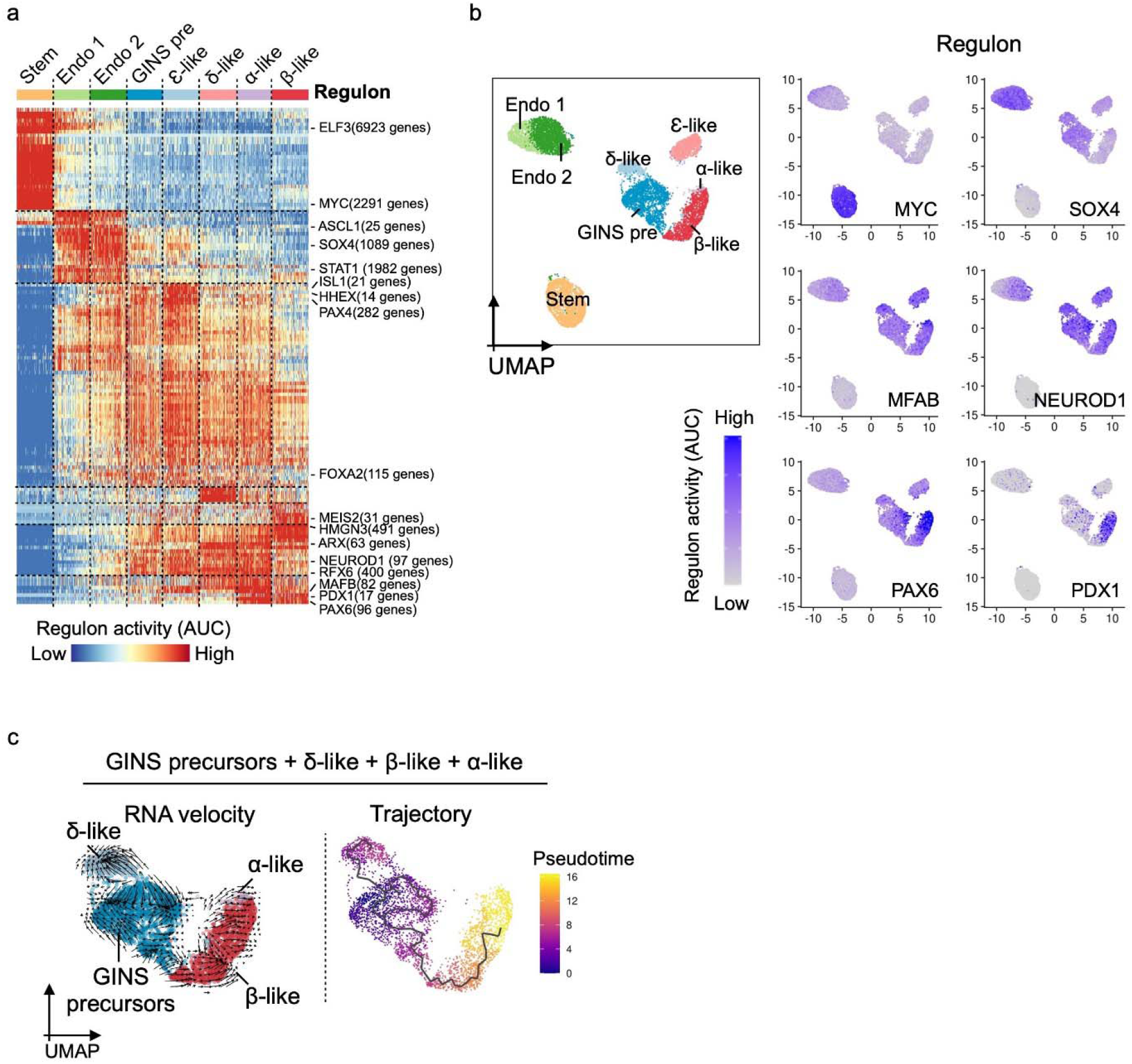
Characterization of the developmental path of GINS organoids. **a,** Heatmap showing waves of transcription factor regulon activations. Key regulons are labeled on the right with the number of their predicted target genes. Stem: hGSCs; Endo 1: endocrine progenitors 1; Endo 2: endocrine progenitors 2; GINS pre: GINS precursors. **b,** Select regulon activity overlaid on UMAP. **c,** RNA velocity and pseudotime trajectory analysis in UMAP showing the developmental path from GINS precursors to endocrine cells in GINS organoids.

**Extended Table 1.**
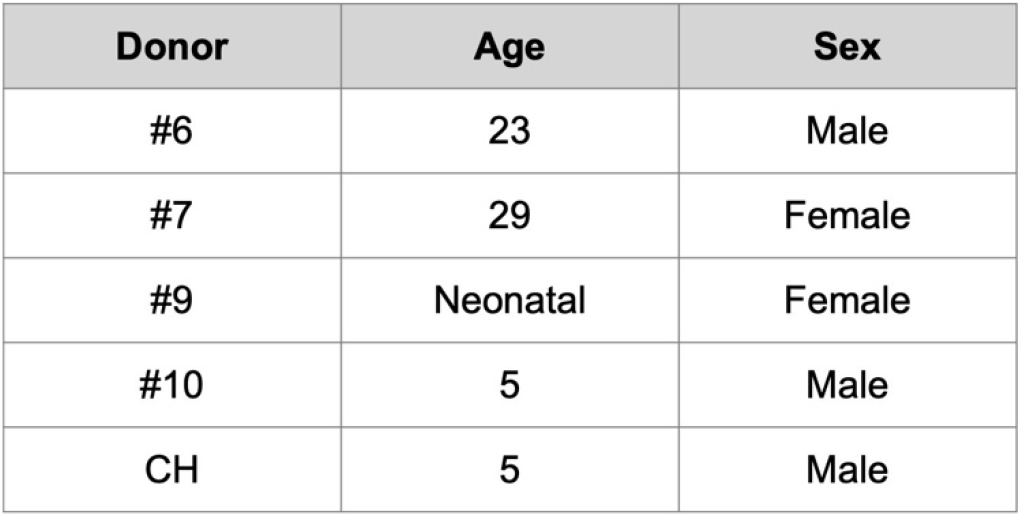
Information of stomach tissue.

| Donor | Age | Sex |
| --- | --- | --- |
| #6 | 23 | Male |
| #7 | 29 | Female |
| #9 | Neonatal | Female |
| #10 | 5 | Male |
| CH | 5 | Male |

**Extended Table 2.** Culture medium and differentiation medium.

| Media | Basal | Supplements |
| --- | --- | --- |
| hGSC medium | 66.7% DMEM<br>33.3% F12K | 18% FBS, 10% R-Spondin-2 conditioned medium, 10 mM Nicotinamide, 1 μM A8301, 5 μg/mL insulin, 10 μM Y-27632, 1 μM DMH1, 50 ng/mL EGF, 2 μM T3 |
| GINS medium | advanced DMEM | 10 mM HEPES, 1X GlutaMAX, 1X B-27, 1X N-2, 500 μM N-Acetyl-L-Cysteine, 10 mM Nicotinamide, 1 μM A8301, and 10 μM Y-27632. |

**Extended Table 3.** Differentiation protocol of GINS organoids.

| Stage | Duration (days) | Medium | Step |
| --- | --- | --- | --- |
| Endocrine progenitors | 2 | hGSC medium | Addition of 1 μM 4-OH-TAM |
| GINS precursors | 2 | 50% hGSC medium<br>50% GINS medium | Lenti-PM infection (MOI 10) |
|  | 2 | 25% hGSC medium<br>75% GINS medium | Medium change |
| GINS organoids | 4-14 | GINS medium | Aggregation and culture |

**Extended Table 4.** Taqman assay list.

| Gene target | Assay number |
| --- | --- |
| *INS* | Hs00355773_m1 |
| *ACTB* | Hs01060665_g1 |
| *GCK* | Hs01564555_m1 |
| *ABCC8* | Hs01093752_m1 |
| *KCNJ11* | Hs00265026_s1 |
| *PDX1* | Hs00236830_m1 |
| *ISL1* | Hs00158126_m1 |
| *NKX6-1* | Hs00232355_m1 |
| *PAX6* | Hs01088114_m1 |
| *NKX2-2* | Hs00159616_m1 |
| *SLC30A8* | Hs00545183_m1 |
| *GCG* | Hs01031536_m1 |
| *GHRL* | Hs01074053_m1 |
| *SST* | Hs00356144_m1 |

## Acknowledgment

We thank Dr. Xi He from Children’s Hospital Boston for the RS2 cell line, Drs. Sean Houghton and David Redmond for initial processing of the scRNA-seq data and for bioinformatic advice, the Starr Foundation Tri-Institutional Stem Cell Derivation Laboratory (Dr. Raphael Lis and Mr. Tyler Lu) and WCM Flow Cytometry Core Facility (Jason McCormick, Tomas Baumgartner) for flow cytometry. We thank Capucine Martin for reviewing the manuscript. We thank WCM CLC Microscopy & Image Analysis Core (Lee Cohen-Gould, Juan Pablo Jimenez) for transmission electron microscopy, and WCM Genomics Resources Core Facility (Jenny Xiang, Xing Wang, Dong Xu) for 10x genomics and next-generation sequencing. We are grateful to Drs. Angie Chi Nok Chong and Shuibing Chen for helping with dynamic GSIS, NDRI and HAM for providing some of the human samples, and Prodo lab for supplying human islets. This work was supported by awards from NIDDK (R01 DK106253, R01 DK13332, R01 DK125817 and UC4DK116280).

## Author Contributions

XH and QZ designed all the experiments, interpreted data, and wrote the manuscript. XH performed all the experiments and the scRNA-seq data analysis. WG participated in all the animal surgery. JZ and YL were involved in stomach sample processing, gastric stem cell isolation, and cell line establishment. JLC collected data from the animal experiments. SL performed independent validation of the functionality of the GINS organoids. WG, JY, CP, JL, HK, and JZ contributed to conducting preliminary experiments. JS produced antibodies against ENTPD3.

## Methods

### Lentivirus packaging and titration

The lentiviruses were packaged as previously described [STAR Protocols (2022), https://doi.org/10.1016/j.xpro.2022.101308]. Viral supernatant was cleared by centrifugation followed by filtration using a 0.45 μm polyethersulfone (PES) filter. Viruses were then concentrated 20-fold by Centrifugal Filter (Millipore Sigma UFC910024). Concentrated viruses were aliquoted and stored at −80°C. To measure the titer of lentiviruses, HEK293FT cells were seeded in 24-well plates to be ~90% confluent at infection. Virus was diluted in DMEM medium containing 10% FBS and 10 μg/mL polybrene and added to the cells. Forty-eight hours post infection, the number of fluorescence-positive cells was counted under a fluorescence microscope if the lentiviral vector carried a fluorescence marker (e.g., mCherry for *Lenti-EF1a-Ngn3ER-mCherry*). Otherwise, transduced cells were visualized by an immunofluorescence assay (e.g., MAFA staining for Lenti-PM). Viral titers were defined by the transduction units per mL (TU/mL).

### Human samples

All studies involving human samples were approved by the ethical committee at Weill Cornell Medical College. Human stomach samples were obtained from Children’s Hospital Boston, the International Institute for the Advancement of Medicine (IIAM), and The National Disease Research Interchange (NDRI). Patient consent was received for these studies. Subject details are described in Extended Table 1. Human islets were obtained from Prodo Labs.

### Derivation and culture of human gastric stem cells from primary mucosal tissues

Human gastric samples (3-5 mm in size) were vigorously washed with cold DPBS 3 times and cut into smaller pieces with a sharp scalpel. Tissue was then incubated with F12K medium containing 2 mg/ml collagenase type IV (Worthington LS004188) at 37°C with pipetting every 5-10 min until most of the glandular cells were released and appeared in solution as clusters. Cells were neutralized with F12K supplemented with 10% FBS and centrifuged at 500 × g for 5 min. Pelleted cells were resuspended in human gastric stem cell culture medium (hGSC medium) and seeded on mitomycin-C-inactivated mouse embryonic fibroblasts (MEF, E13.5-14.5, DR4 strain, The Jackson Laboratory 003208) coated 3-cm dish.

The hGSC medium (Extended Table 2) was formulated using a method slightly modified from a published report^9^: basal medium composed of 66.7% DMEM (high glucose, Thermo Fisher Scientific, 11-965-118) and 33.3% F12K (Thermo Fisher Scientific, 21127030) was supplemented with 18% FBS (R&D systems, S11150), 10% R-Spondin-2 conditioned medium, 10 mM Nicotinamide (Sigma-Aldrich, N5535), 25 μM Primocin (Invivogen, ant-pm-1), 1 μM A8301 (Cayman, 9001799), 5 μg/mL insulin (Sigma-Aldrich, 10516-5ML), 10 μM Y-27632 (LC Laboratories, Y-5301), 1 μM DMH1 (Cayman, 16679), 50 ng/mL EGF (R&D Systems, 236-EG-01M), and 2 μM T3 (Sigma-Aldrich, T6397).

hGSCs were maintained at 37 °C in a 7.5% CO_2_ incubator. Culture medium was changed every 2-3 days. Y-27632 was withdrawn 24 h post passage. hGSC colonies were split every 4-6 days at a ratio between 1:3 and 1:5 as follows: cells were washed twice with DPBS and dissociated by 10-12 min incubation in TrypLE (Thermo Fisher Scientific, 12604021) with pipetting at the end. Cells were then neutralized by DMEM medium with 10% FBS and centrifuged at 300 x g for 5 min. Pelleted cells were resuspended in hGSC medium and then seeded on an inactivated-MEF-coated dish.

### Human gastric stem cell engineering with Lentiviral infection for transgene and reporter gene expression

To engineer hGSCs, cells were passaged in one well of 6-well plate 24 hours prior to lentiviral transduction. Cells were washed with DPBS and overlaid with hGSC medium containing 10 μg/mL polybrene and 25 μL of lentivirus (viral titer: ~10^8^TU/mL). Spinfection was then performed as follows: the cell culture with lentivirus was spun at 1000 × g for 30 min at 37°C and then incubated at 37°C in a 7.5% CO_2_ incubator for 48 hours. The medium was changed to hGSC medium containing 1 μg/mL puromycin or 10 μg/mL Blasticidin according to the selection marker incorporated into the construct for 2 weeks. The *Ngn3ER*-hGSCs were labeled with constitutive mCherry expression by incorporation of a polycistronic cassette EF1α-*Ngn3ER-Puro^R^-mCherry* (*Puro^R^*, puromycin resistant gene). To establish the Ngn3 and Pdx1-Mafa dual inducible cell line (*Ngn3ER/PM-hGSCs*), *Ngn3ER-hGSCs* were infected by lentivirus carrying polycistronic cassettes *TetO-Pdx1-Mafa and* PGK-*rtTA-Blast^R^* (*Blast^R^*, blasticidin resistant gene). To establish the pGAL-*GFP* reporter cell line (*Ngn3ER*/pGAL-*GFP*-hGSCs), *Ngn3ER*-hGSCs were infected by lentivirus carrying transgenic *GAL* promoter (1,591 bp, upstream region of the TSS (transcription start site)) driven GFP reporter (pGAL-*GFP*) and *PGK-Blast^R^*. Doxycycline (Dox, 1 μg/mL) or 4-OH-Tam (1 μM, Sigma-Aldrich, H7904) were added to the medium to induce the expression of TetO-driven transcription or Ngn3ER activation.

### Supplement screen to formulate chemically defined serum-free medium for GINS differentiation

*Ngn3ER/PM*-hGSCs were seeded 5 days prior to differentiation in 96-well plate. To start differentiation, 1 μM 4-OH-TAM was added and incubated for 2 days. The culture medium was then changed to serum-free basal medium with one supplement and Doxycycline to activate *Pdx1-Mafa* expression. The basal serum-free medium for screens was prepared as follows: advanced DMEM/F12 (Thermo Fisher Scientific, 12634010) was supplemented with 10 mM HEPES (Thermo Fisher Scientific, 15-630-080), 1X GlutaMAX (Thermo Fisher Scientific, 35050061), 1X B-27 (Themo Fisher Scientific, 17504044), 1X N-2 (Thermo Fisher Scientific, 17502048), 25 μM Primocin, and 500 μM N-Acetyl-L-Cysteine (NAC) (Sigma-Aldrich, A9165). Seven days after supplement treatment, we assessed (1) spontaneous clustering of nascent GINS cells by live imaging of mCherry^+^ cells, and (2) qPCR of *INS* mRNA levels. We then analyzed multiple β-cell markers with qPCR on samples treated with Nicotinamide, Y-27632 and A8301.

### Generation of GINS organoids (Extended Table 2–3)

1. <u>Ngn3ER activation (Differentiation to endocrine progenitors) (day 0-2):</u> *Ngn3ER*-hGSCs were seeded 4-5 days prior to differentiation. Cells were washed with DPBS and overlaid with hGSC medium containing 1 μM 4-OH-TAM.
2. <u>Pdx1-Mafa transduction (Differentiation to GINS precursors) (day 2-6):</u> Endocrine progenitors were gently washed with DPBS, incubated in DPBS for 5-10 min, and detached by pipetting. Pelleted cells were then digested in TrypLE at 37°C for 10 min with pipetting every 3-5 min. Dissociated endocrine progenitors were transduced by Lenti-PM at an MOI of 10 by spinfection in medium composed of 50% of hGSC medium, 50% of GINS medium, and 10 μg/mL polybrene. Cells were then transferred to tissue culture dishes (~10^7^ cells per 10-cm dish) coated with Fibronectin (1:50, Sigma-Aldrich, F4759) and Matrigel (1:50, VWR, 47743-722). On day 4, the culture medium was changed to medium consisting of 75% GINS medium and 25% hGSC medium.
3. <u>GINS organoid formation (day 6-21):</u> GINS precursors were dissociated by 5-10 min TrypLE treatment and aggregated (typically 2.0-2.4 million cells/well) in AggreWell400 (STEMCELL Technologies, 34450) using the manufacturer’s recommended protocol. Medium was changed every 2-3 days. Aggregates normally formed within 24 hours.
4. <u>GINS medium for this study was formulated as the follows:</u> advanced DMEM/F12 supplemented with 10 mM HEPES, 1X GlutaMAX, 25 μM Primocin, 500 μM NAC, 1X B-27, 1X N-2, 10 mM Nicotinamide, 1 μM A8301, and 10 μM Y-27632.

### Static glucose-stimulated insulin secretion (GSIS)

Ten to twenty GINS organoids were sampled for each group. Organoids were washed with RPMI-1640 no nutrient medium (RPMIN, MyBioSource, MBS652918), and equilibrated in 3.3 mM glucose RPMIN for 2 hours. Organoids were incubated in low-glucose RPMIN for 1 hour, and supernatant was collected. Organoids were then incubated in high-glucose RPMIN for 1 hour, and supernatant was collected. For sequential glucose challenge, organoids were washed 2 times in RPMIN prior to the next stimulation. Secreted insulin was measured using the Stellux Chemi Human Insulin ELISA (ALPCO Diagnostics, 80-INSHU-CH10).

### Dynamic GSIS with perifusion

Twenty GINS organoids were sampled for each group. Organoids were washed with RPMIN and equilibrated in 1 mM glucose RPMIN for 2 hours at 37°C incubator. The assay was performed in RPMIN on a temperature-controlled (37°C) perifusion system (Biorep Technologies). Organoids were loaded in chambers and perifused at a flow rate of 100 μL/min in the following steps: 48 min in 1 mM glucose, (2) 24 min in 2 mM glucose, (3) 36 min in 20 mM glucose, (4) 24 min in 20 mM glucose with 100 ng/mL Liraglutide (Cayman, 24727), (5) 48 min in 2 mM glucose, (6) 20 min in 2 mM glucose with 30 mM KCl. Secreted insulin was measured using the Stellux Chemi Human Insulin ELISA.

### Transplantation studies in normoglycemic and hyperglycemic mice

All mouse experiments were conducted under the IACUC protocol 2018-0050 at Weill Cornell Medical College. Mice were housed in a temperature- and humidity-controlled environment with 12 hours light/dark cycle and food/water ad libitum. Transplantations were performed with male NSG mice. GINS organoids or human islets were transplanted under the capsule of the left kidney in NSG mice anesthetized with isoflurane. For glucose tolerance test, transplanted mice were fasted for 6 hours and injected with 2 g/kg glucose intraperitoneally. Blood glucose was then measured at indicated time points by glucometer. For in vivo GSIS, transplanted mice were fasted overnight and injected with 2 g/kg glucose intraperitoneally. Blood before injection and 1 hour post glucose injection was collected by submandibular bleeding with Microvette 300 capillary blood collection tube (Sarstedt, 20.1309.100). Serum was separated from the blood for insulin measurement by Stellux Chemi Human Insulin ELISA. For the diabetes rescue experiment, male mice were injected with four doses of Streptozotocin (35 mg/kg/d) on four consecutive days to induce hyperglycemia. Mice that showed hyperglycemia (>250 mg/dl) on four consecutive days were selected for transplantation. Sham transplantation was conducted by operating the surgical procedure without infusing cells. Random fed blood glucose was monitored 2 times per week. To remove grafts, a survival nephrectomy was performed after 90-100 days post transplantation. Briefly, the left kidney was ligated at the renal hilum using 3-0 silk and then resected.

### Immunofluorescence

GINS organoids were fixed in 4% PFA for 15 min at room temperature. Kidney samples were fixed in 4% PFA at 4°C for 1 hour. Samples were washed in PBS, incubated in PBS containing 30% sucrose overnight. Samples were frozen in OCT (Tissue-Tek) next day, and then cryosectioned. Following PBS wash, sections were blocked for 1 hour at room temperature in blocking buffer: 10% normal donkey serum (Jackson ImmunoResearch, 017-000-121) in PBST (0.1% TritonX-100 in PBS). Sections were then incubated with primary antibodies in blocking buffer overnight at 4°C. The following primary antibodies were used in this study: rat anti-C-peptide (DSHB, GN-ID4), guinea pig antiinsulin (Dako, A0564), goat anti-ghrelin (Santa Cruz, sc-10368), guinea pig anti-glucagon (Linco, 4031-OlfZ), rabbit anti-somatostatin (Dako, A0566), rabbit anti-MAFA (Bethyl, A700-067), rabbit anti-PAX6 (Millipore, AB2237), mouse anti-NKX2-2 (DSHB, 74.5A5-s), rat anti-CD31 (BD Pharmingen, 550274), mouse anti-ENTPD3 (developed in house), rabbit anti-PCSK1 (Millipore, AB10553), and mouse anti-GAL (Santa Cruz, sc-166431). Slides were washed three times in PBST, followed by secondary antibody incubation in blocking buffer with DAPI for 1 hour at room temperature (protected from light). Following 3 washes in PBST, slides were mounted in mounting medium (Vector Laboratories, H-1700-10) and covered with coverslips. The representative images were captured using either a confocal microscope (710 Meta) or a Nikon fluorescence microscope.

### RNA extraction, reverse transcription, and real-time PCR

RNA was extracted (Qiagen, 74034) and reversely transcribed (Thermo Fisher Scientific, 43-688-13) to complementary DNA (cDNA). cDNA was diluted and then quantified by real-time PCR with TaqMan assay listed in Extended Table 4. For some experiments, Cells-to-Ct Kit was used (Thermo Fisher Scientific, A35377).

### Fluorescence-activated Cell Sorting (FACS)

For quantitative flow cytometry, GINS organoids were dissociated in TrypLE for 40 min. Dissociated cells were stained with Fixable Viability Dye 455UV (Thermo Scientific, 65-0868-14) according to the manufacturer’s manual. Cells were then fixed and permeabilized using Intracellular Fixation & Permeabilization Buffer Set (Thermo Scientific, 88-8824-00) according to the manufacturer’s manual. Fixed cells were then incubated in 1X permeabilization buffer with primary antibodies for 1 h at room temperature and washed with permeabilization buffer for 3 times and resuspended in flow cytometry staining buffer (Thermo Scientific, 00-4222). Stained cells were then passed through a 40 μm nylon strainer before being sorted (Supplementary Fig. 1). For live cells sorting, cells were dissociated in TrypLE (40 min for hGSCs, 5 min for endocrine progenitors, 5 min for GINS precursors and 40 min for GINS organoids), and pelleted. Cells pellet was resuspended in FACS buffer (1% glucose, 10 mM HEPES, 10 μM Y-27632, 1 mM N-acetyl-l-cysteine, and 2% FBS in DPBS) and passed through a 40 μm nylon strainer before being sorted (Supplementary Fig. 2-4).

To purify GAL-GFP^+^ cells, Ngn3ER/pGAL-GFP-hGSCs were seeded and differentiated toward GINS cells. On day 7 post differentiation, cells were sorted for GFP^+^ cells. Sorted cells were then aggregated into organoids and analyzed 1 and 14 days post aggregation.

### Single cell RNA-seq (scRNA-seq) from cultured and transplanted GINS organoids and multiplex scRNA-seq

For the time-course study of GINS generation, samples at different time points were harvested on the same day from parallel cultures. Cells were dissociated in TrypLE. FACS sorting was conducted to purify mCherry^+^ DAPI^-^ cells. Purified samples were then multiplexed according to the 10X genomics protocol CG000391. Briefly, 2×10^5^ sorted cells from each time point were washed in PBS with 0.04% BSA (Millipore Sigma, A1595), and then labeled with multiplexing oligo individually for 5 min at room temperature. Following two washes in PBS with 1% BSA, equal number of labeled cells from different time points were then pooled. In total 30,000 cells were then loaded for 10X genomics. To compare corpus and antral GINS organoids, cells were dissociated by TrypLE on day 21 post induction. To harvest GINS cells from the grafts, the grafts under the kidney capsule were removed with a scalpel and minced. The tissues were digested with type III collagenase (300U/ml in RPMI 1640) for 1 hour, followed by 5-10 min TrypLE treatment with pipetting. Digested tissue was filtered through 40 μm nylon strains and purified by FACS with mCherry^+^ and DAPI^-^ gating. Samples were kept in GINS medium on ice until ready to be processed by 10X genomics single-cell droplet sample preparation workflow at the Genomics Core Facility at Weill Cornell Medicine as previously described^44^.

### scRNA-seq analysis

1. <u>Demultiplexing and reads alignment:</u> Human islets (donor #1, #2, #3, #4 and #9) scRNA-seq datasets were downloaded from GEO database (GSE114297)^29^. Multiplexed sequencing data from the Illumina NovaSeq6000 were demultiplexed using the ‘multi’ pipeline from Cell Ranger (v6.1.2). Each of the four Cell Multiplex Oligo labels was assigned a unique sample ID. All the datasets were then processed with the 10X built mouse and human reference ‘refdata-gex-GRCh38-and-mm10-2020-A’z’.
2. <u>Quality control and count normalization:</u> Cell Ranger outputs were used as input to create Seurat objects by Seurat (v4.0.1)^45^. Cells that express more murine genes than human genes were defined as contaminant murine cells (e.g., murine host cells from kidney) and removed from the datasets. Murine features were then removed. Low-quality cells were removed as follows: In general, cells were considered low-quality if the number of detectable genes or read counts is below the 3^rd^ percentile or above the 97^th^ percentile of the datasets, or percentage of mitochondrial genes is more than 18. *NormalizeData* function was used for normalization with default parameters. Putative doublet cells were identified by DoubletFinder and removed^46^.
3. <u>Scaling, dimension reduction, cell clustering, differential expression analysis and cell annotation:</u> The normalized data was scaled by *ScaleData* function with mitochondrial genes percentage regressed out. Cell cycle status was inferred by *CellCycleScoring* function and regressed out for the time course study. Principal component analysis (PCA) was performed on the scaled data by *runPCA*. To place similar cells together in 2-dimensional space, selected top principal components (PCs) of human islet, GINS organoid (corpus), antral GINS organoid, GINS generation time course and GINS graft (corpus) were used as input respectively in non-linear dimensional reduction techniques including tSNE and UMAP. The same PCs of each sample were used to construct K-nearest neighbor (KNN) graph by *FindNeighbors* function. To cluster cells, *FindClusters* function was used with a range of resolution between 0.2 to 2. Cells clustered by different resolutions were all visualized by *DimPlot* function. To identify markers of each cell cluster, *FindAllMarkers* function was implemented using the following parameters: only.pos = TRUE, min.pct = 0.3, logfc.threshold = 0.3. Human islets cells were firstly automatically annotated by the R package SingleR with a publish islets dataset as reference ^31^. The annotation of the islet dataset was then slightly modified according to the clustering results and cluster markers. Non-endocrine cells or unclear cell types were removed from the human islet dataset. All the other datasets were annotated manually according to markers and integration results.
4. <u>Integration analysis:</u> GINS organoids (corpus) were integrated with antral GINS organoids, human islets endocrine cells, and GINS grafted cells respectively. The genes that were used for integration were chosen by *SelectIntegrationFeatures* function with default parameter. Integration anchors were identified by *FindIntegrationAnchors* function and used to integrate two datasets together with *IntegrateData* function. All the integrated datasets were scaled with mitochondrial genes regressed out. Dimensions were reduced by PCA, tSNE and UMAP.
5. <u>Identity scoring:</u> Signature gene sets (Supplemental Table 3) were downloaded from cell type signature gene sets (C8 collection) of Molecular Signature Database (v7.5.1). Specifically, the gastric signature is a gene list containing curated cluster markers for gastric chief, immature pit, mature pit, isthmus, neck, and parietal cells identified in the study^32^, while the signature of β-cell is the MURARO_PANCREAS_BETA_CELL gene set^31^. Both gene sets were then applied as inputs in the *AddModuleScore* function of Seurat with default parameters, which calculates module scores for feature expression programs on single-cell level.
6. <u>Pseudotime trajectory, RNA velocity, regulon and Gene Oncology (GO) analysis:</u> Seurat object of the time course dataset was converted to monocle 2 CellDataSet object^47–49^. Size factor and dispersion were estimated by *estimateSizeFactors* and *estimateDispersions* function respectively. Cell type markers identified by Seurat were sorted based on adjusted p-value. Top 100 markers of each cell type were marked by *setOrderingFilter* function for later trajectory construction. Data dimension was then reduced by *reduceDimsion* function with the following arguments: max_components = 2, method = ‘DDRTree’)-Cells were then ordered along the trajectory by *orderCells* function. To build pseudotime trajectory on UMAP, Seurat object was converted to monocle 3 object by *as.cell_data_set* function. Cells were then clustered by *cluster_cells* function. The trajectory of GINS precursor, δ-like, β-like and α-like cells, which were in the same partition, was constructed by *learn_graph* followed by *order_cells* function. RNA velocity was evaluated by *RunVelocity* function provided by R packages velocyto.R and SeuratWrappers^50^. Regulon activity was computed by SCENIC R package as previously described^51^. GO analysis was done by *enrichGO* function in ClusterProfiler as previously described^52,53^.

## Statistical analyses

Statistical analyses were done using GraphPad Prism 9 or R 4.0.5. P values, sample size, and statistical methods are described in figure legends.

## Data availability

Data is available upon request.

## Code availability

Code is available upon request.

