## Supplemental figures for "Restoring glucose homeostasis with stomach-derived human insulin-secreting organoids"

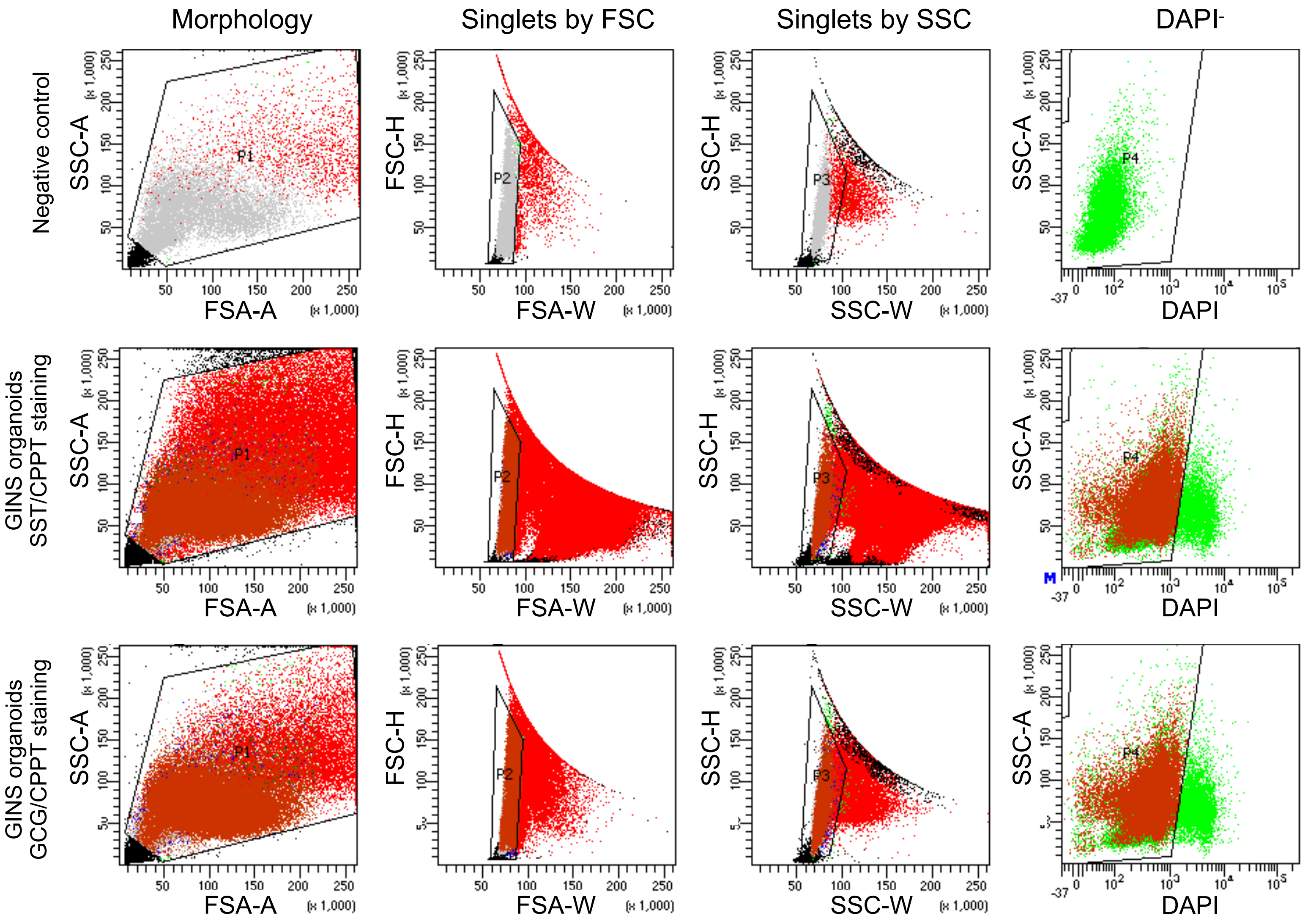


**Supplementary Fig. 1 | Flow cytometry gating strategy for GINS organoids.**

a

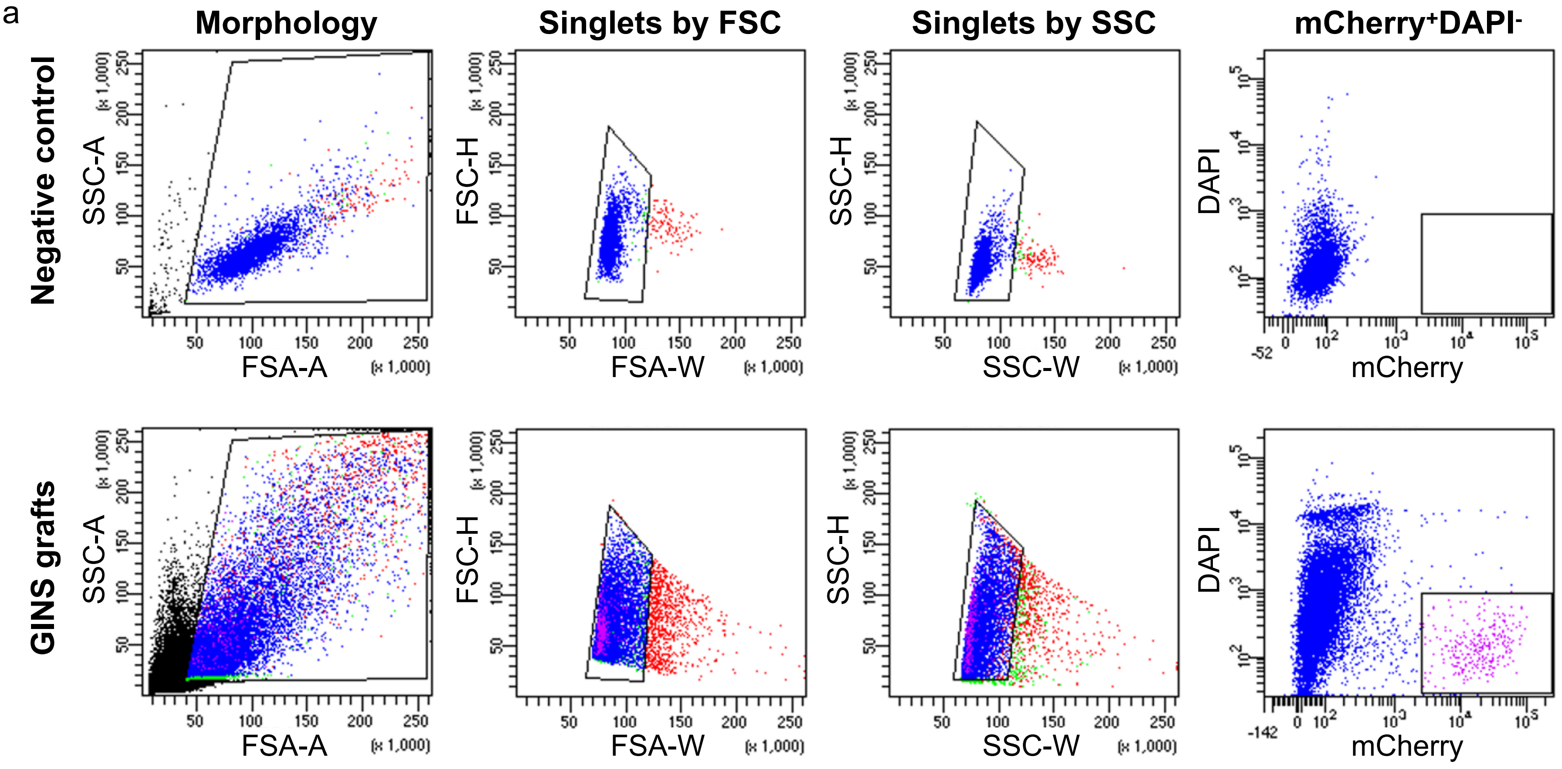


b GINS grafts statistics

| Population | %Parent | %Total |
|---|---|---|
| All Events | #### | 100.0 |
| Morphology | 22.2 | 22.2 |
| Singlets FSC | 93.8 | 20.8 |
| Singlets SSC | 89.7 | 18.7 |
| DAPI- mCherry+ | 2.1 | 0.4 |

**Supplementary Fig. 2 | Flow cytometry gating strategy for GINS grafts. a**, Sorting for mCherry+DAPI- cells. **b**, Percentage of cell populations.

a

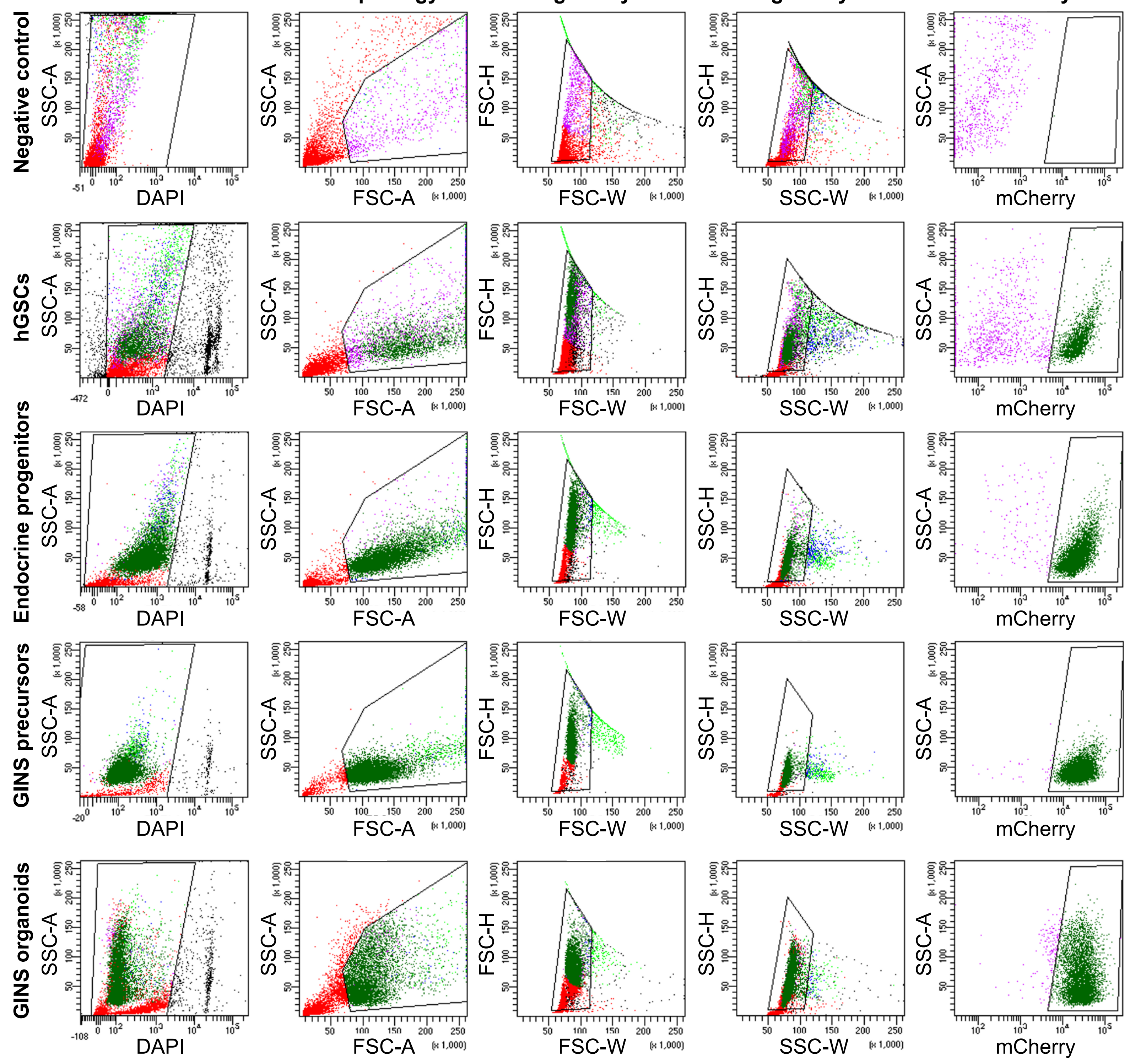


b

**hGSC**

| Population | %Parent | %Total |
|---|---|---|
| All Events | #### | 100.0 |
| Live | 77.4 | 77.4 |
| Morphology | 47.3 | 36.6 |
| Singlet FSC | 83.9 | 30.7 |
| Singlet SSC | 92.7 | 28.5 |
| mCherry+ | 67.6 | 19.3 |

**Endocrine progenitors**

| Population | %Parent | %Total |
|---|---|---|
| All Events | #### | 100.0 |
| Live | 94.8 | 94.8 |
| Morphology | 73.9 | 70.0 |
| Singlet FSC | 95.1 | 66.6 |
| Singlet SSC | 96.8 | 64.4 |
| mCherry+ | 98.0 | 63.1 |

**GINS precursors**

| Population | %Parent | %Total |
|---|---|---|
| All Events | #### | 100.0 |
| Live | 98.0 | 98.0 |
| Morphology | 91.7 | 89.9 |
| Singlet FSC | 96.5 | 86.8 |
| Singlet SSC | 99.0 | 85.9 |
| mCherry+ | 99.7 | 85.6 |

**GINS organoids**

| Population | %Parent | %Total |
|---|---|---|
| All Events | #### | 100.0 |
| Live | 94.1 | 94.1 |
| Morphology | 56.2 | 52.9 |
| Singlet FSC | 97.0 | 51.3 |
| Singlet SSC | 99.1 | 50.8 |
| mCherry+ | 97.2 | 49.4 |

**Supplementary Fig. 3 | Flow cytometry gating strategy for hGSCs, endocrine progenitors, GINS precursors and GINS organoids. a**, Sorting for $mCherry^{+}DAPI^{-}$ cells. **b**, Percentage of cell populations.

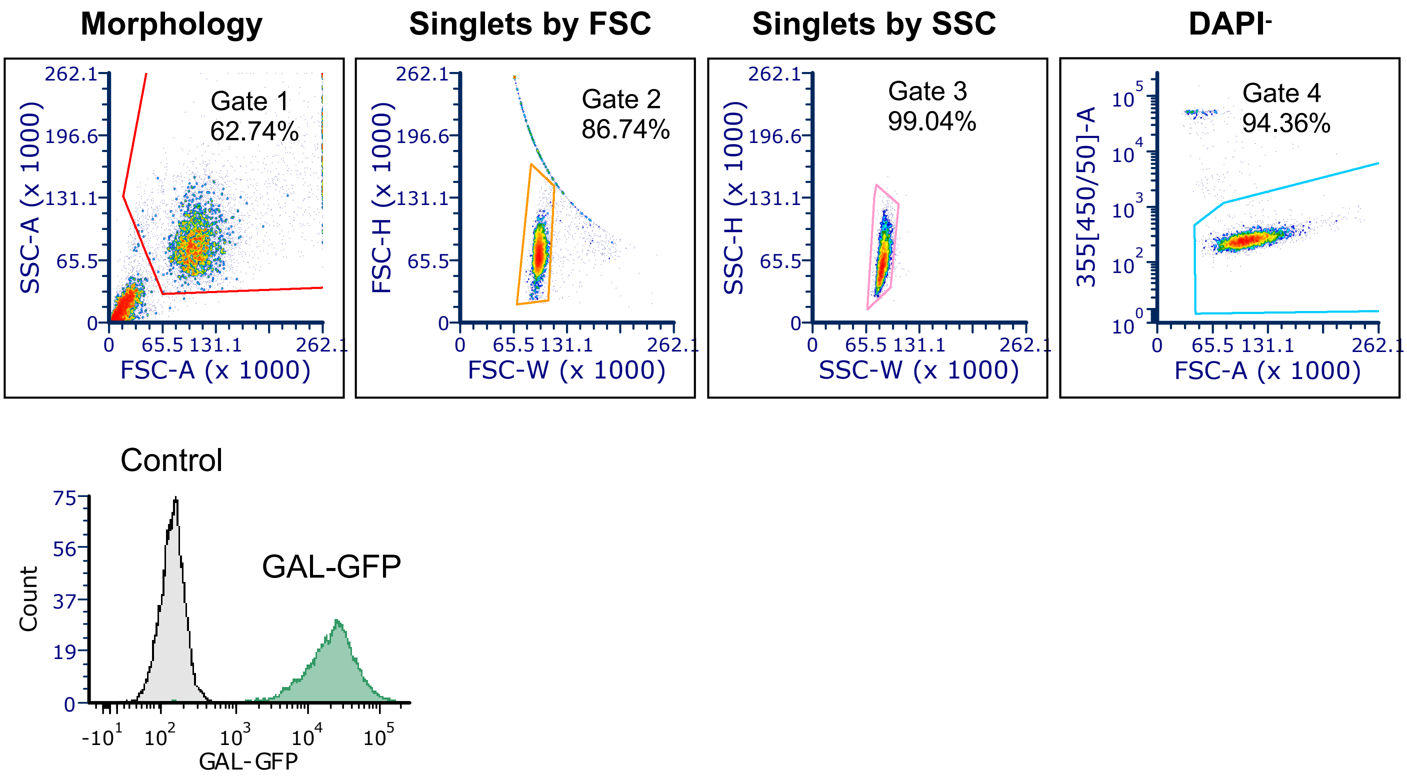


**Supplementary Fig. 4 | Flow cytometry gating strategy for GINS organoids with GAL-GFP reporter.**
